# DOPA residues endow collagen with radical scavenging capacity

**DOI:** 10.1101/2023.01.23.524231

**Authors:** Markus Kurth, Uladzimir Barayeu, Hassan Gharibi, Andrei Kuzhelev, Kai Riedmiller, Jennifer Zilke, Kasimir Noack, Vasyl Denysenkov, Reinhard Kappl, Thomas F. Prisner, Roman A. Zubarev, Tobias P. Dick, Frauke Gräter

## Abstract

Here we uncover collagen, the main structural protein of all connective tissues, as a redox-active material. We identify dihydroxyphenylalanine (DOPA) residues, post-translational oxidation products of tyrosine residues, to be common in collagen derived from different connective tissues. We observe that these DOPA residues endow collagen with substantial radical scavenging capacity. When reducing radicals, DOPA residues work as redox relay: they convert to the quinone and generate hydrogen peroxide. In this dual function, DOPA outcompetes its amino acid precursors and ascorbic acid. Our results establish DOPA residues as redox-active side chains of collagens, probably protecting connective tissues against radicals formed under mechanical stress and/or inflammation.

## Introduction

Protection mechanisms against free radicals are vital to any living organism. Radicals can cause tissue damage, are associated with disease, and accelerate aging (*1–4*). To prevent damage by nonspecific reactions with proteins, lipids and nucleic acids, nature has come up with highly efficient radical scavengers which react with and stabilize radicals. A prominent example is ascorbic acid (*5*).

The catechols dihydroxyphenylalanine (DOPA) and dopamine have various biological functions independent from radical biology, but one important aspect also for them is their excellent radical scavenging capacity (*6, 7*). This property is based on the high thermodynamic stability of the semiquinone radical, the intermediate species between the catechol and the quinone upon successive oxidation (Fig. 1) (*7*). It is harnessed in vivo, e.g. by DOPA-containing proteins in mitochondria to capture superoxide from oxygen-leakage (*8, 9*), or by melanin in skin to capture radicals from light (*10, 11*), and also exploited technologically, among others in form of polydopamine as a mimic of melanin and efficient antioxidant.

**Figure 1.**
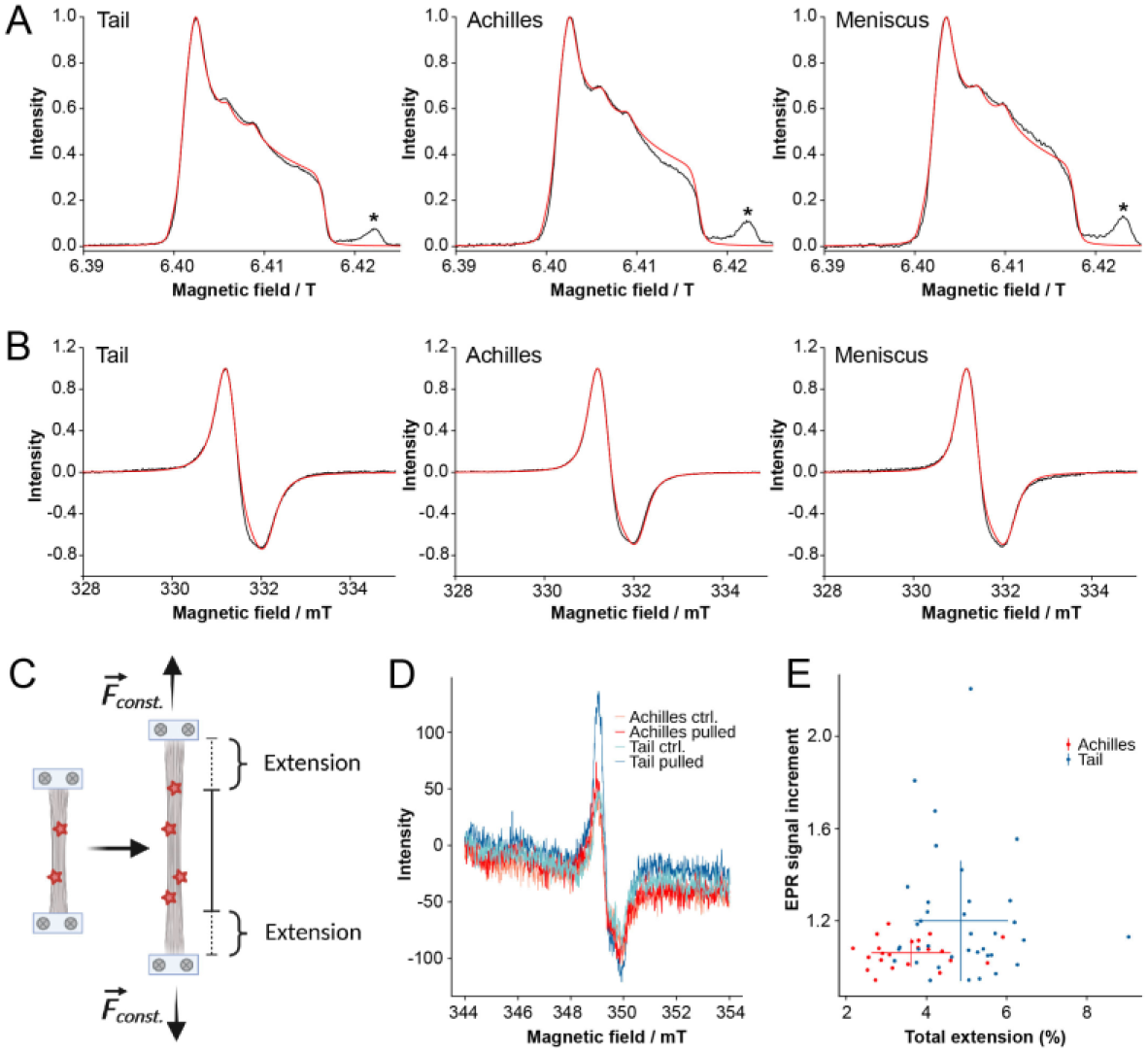
DOPA mechanoradicals are formed in different collagen type I tissues. (**A**) G-band EPR spectra measured at 40 K and (**B**) X-band EPR spectra measured at 298 K for ground collagen tissue (Tail, Achilles’, meniscus, samples I, black curves) showed a similar radical signature (each curve represents one biological replicate, n=2). Fitting with EasySpin (*20*) (simulation shown as red curves) showed DOPA anion radical (gx=2.0071, gy=2.0066, gz=2.0022) as well as two unknown organic radicals (gx=2.0075, gy=2.0055, gz=2.0021 and gx=2.0075, gy=2.0046, gz=2.0021). The signal marked by * in the G-band spectra in (A) appears from the quartz capillary. (**C**) In the pulling experiments, tendons were fixed at both sides and pulled at constant force (here: 15 gmf or ~15 N) on an extensometer which also recorded the stress strain curve. From this curve, the extension was calculated (see Supplementary Fig. 1). Created with BioRender.com. (**D**) Representative spectra of a X-band EPR measurement of tail and Achilles’ tendon tissue before (control/Ctrl.) and after mechanical stretching at 15 N for 1000 sec (pulled). The EPR increment was calculated by dividing the peak difference maximum to minimum of both measurements (Pulled/Ctrl.) and (**E**) plotted against the total extension calculated from the strain curve.

Radicals can virtually occur in any tissue (*12*). Capturing them on site requires a high local abundance of potent radical scavenging molecules, and catechol containing protein materials would be ideal candidates for this purpose. If any protein that is ubiquitously and substantially expressed across tissues has a general built-in function as radical scavenger and cellular protectant is unknown.

Collagen I is the most abundant protein of our body, and makes up the larger part of our connective tissue (*13*). It has been long thought to act as a scaffolding and force-bearing protein with primarily mechanical functions, and a direct role in radical biochemistry has remained unrecognized. Uncommon types of collagen that naturally contain DOPA, however, are well known, namely in mussel byssal threads (*14*). There, DOPA chelates metal ions and thereby equips collagen with an adhesive function (*15*), an observation which served as a major inspiration for using catechol chemistry in synthetic polymer adhesives. More recently, we obtained first incidental evidence for naturally occurring DOPA in collagen I, the major collagen of connective tissue, namely at one specific tyrosine site (Tyr1014DOPA in α(II)) in rat tail tendon (*16*). Our findings hinted towards a function of DOPA as a scavenger of mechanoradicals (*16*).

If collagen I generally contains DOPA and can serve as a radical scavenger, as a common theme across tissues, is unknown. Interestingly, exposing collagen I to UV light can artificially cause the conversion of tyrosines into DOPA (*17*),and enzymatic pathways of tyrosine oxidation to DOPA exist, too (*18*). Also, tyrosines (and phenylalanines) are conserved in collagen I (*16*), and their role as DOPAs could explain this evolutionary constraint.

Here, we show that DOPA naturally occurs in collagen type I polypeptides and functions as a radical scavenger in tail, Achilles’ tendons and meniscus. DOPA’s catechol moiety very efficiently stabilizes mechanoradicals and generates hydrogen peroxide in the degradation process, in this capacity outcompeting ascorbic acid.

## Results and Discussion

### DOPA radicals form in different collagen type I tissues

Previously, we had identified DOPA phenoxy type radicals in tail tissue of Rattus norvegicus as the most stable mechanoradical type (*16*). To find out if that radical type is also occurring in other collagen type I-rich tissues, we mechanically stressed tail, Achilles’ and meniscus tissues. The occurring radicals were detected in the tissue by G-band (180 GHz) electron paramagnetic resonance (EPR) spectrometry at 60 K (Figure 1A, Supplementary Figure S1A). The fitted EPR spectra showed that the majority of signal in studied samples corresponded to a DOPA anion radical (g_x_= 2.0071, g_y_= 2.0066, g_z_= 2.0022; ≥ 66% for tail, ≥ 57% for Achilles’, ≥ 54% for meniscus), while the rest probably corresponds to two further, yet unknown organic radicals with small hyperfine interactions (Supplementary Figure S2). For continuous wave (cw) X-band EPR measurements, the shape of the singlet signals was almost indistinguishable in all tissues (Figure 1B, Supplementary Figure S1B). Overall, DOPA appeared as the chemical function carrying most radicals generated in stressed collagen I tissues (Supplementary Table S1). Hence, we were able to confirm the role of the DOPA anion as the most efficient radical scavenger, as we previously reported for mechanoradicals in tail tendon (*16*).

Next, we aimed to quantify the formation of mechanoradicals after stretching of tendon tissue; due to the small size, no such measurements were done for meniscus tissue. For continuous wave (cw) X-band EPR measurements, we stretched tail or Achilles’ tendons. In the extensometer, the samples were pulled for 1000 s at a constant force of 15 N, which corresponds to approximately 10-15 MPa (tissue section of ~1.0 - 1.5 mm^2^) (Figure 1B). The analysis of the stress strain curve showed that the Achilles tendons were on average significantly stiffer with a mean extension of 3.6 ± 1.0% (mean ± standard deviation) then the tail tendons with an extension of 4.9 ± 1.1% (Mann-Whitney, two-tailed, p<0.0001) (see also Supplementary Fig. S1E+F). This we attribute to the different crosslinking pathways in these two tissues (*19*). For each tendon sample, we recorded the amount of radicals before and after the stretching experiment by quantifying peak height of the singlet single. Before stretching, there was always a basic signal of (mechano-)radicals in the tissue which we attribute to the preparation and drying of the tissue (Figure 1C). After stretching, the signal in the EPR had increased just marginally for the Achilles’ tendon by a factor of 1.06 ± 0.06 while this factor was significantly higher for the tail tendon with 1.20 ± 1.27 (Mann-Whitney, two-tailed, p=0.0227) (Supplementary Fig. S1C-F). Also, the recorded signals in tail tendons were overall stronger as compared to the Achilles’ tendons. When correlating the increment of radicals after stress with the extension of tissue, we found that the tail tendons were bent more easily when pulled at a constant force and at the same time showed a larger increment of the radical signal as compared to the stiffer Achilles tendons (Figure 1D).

### DOPA is a modification by oxidation of tyrosine residues

Having identified DOPA as anion radical in tissue, we wanted to test its occurrence in the tissues as stable post-translational modification. Using a mass spectrometric (MS) based peptide sequencing approach, we determined the fraction of DOPA in the collagen type I tissues that had shown a DOPA radical signature after mechanical stress (Figure 1+2). Here, DOPA may originate from a single oxidation event of a tyrosine (Tyr) residue or a double oxidation of a phenylalanine (Phe) residue (*21*) (Figure 2A).

**Figure 2.**
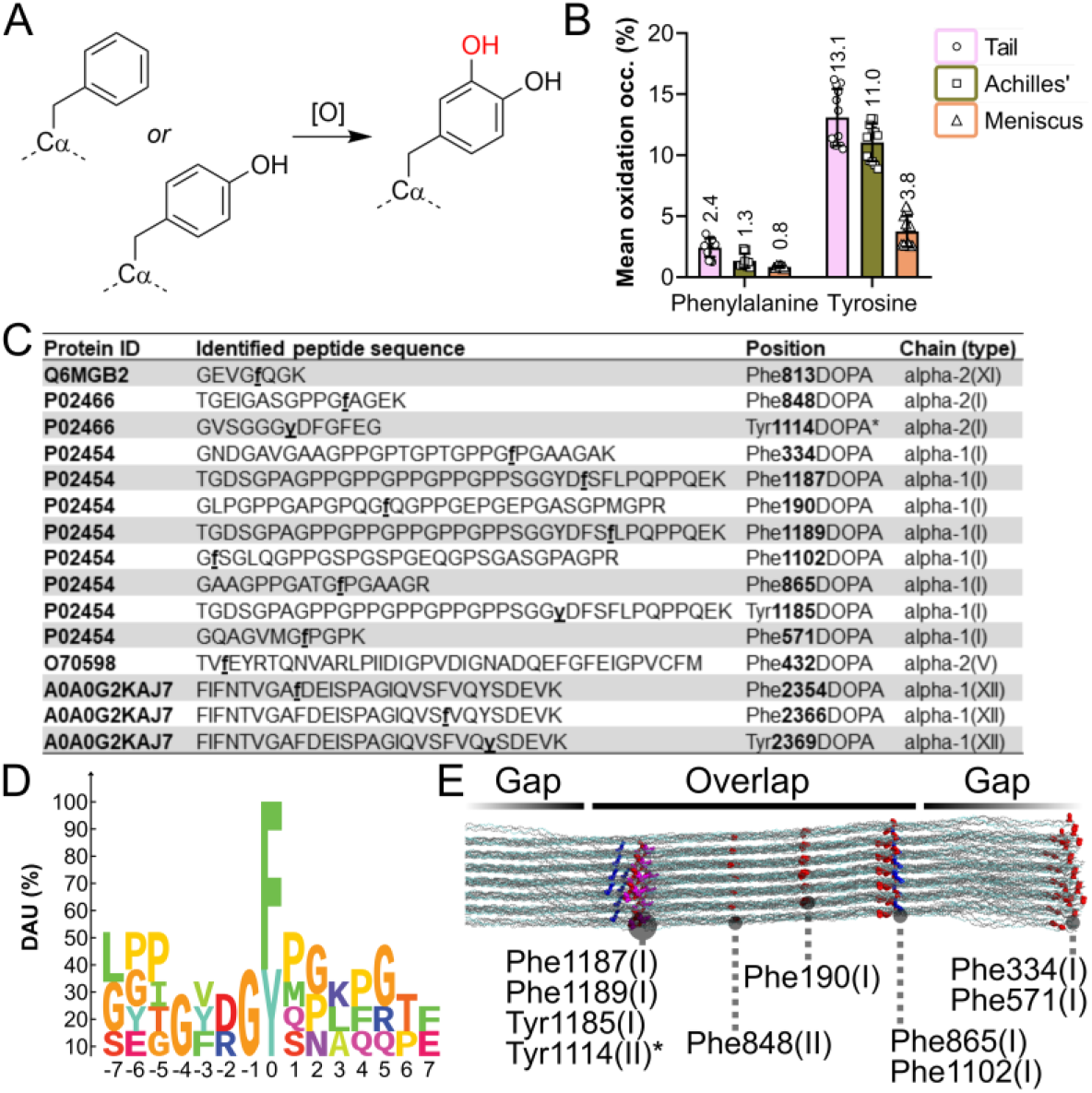
DOPA is a modification by oxidation of phenylalanine or tyrosine residues. (**A**) Both Phe and Tyr residues can be double or single oxidized [O], respectively, to generate a DOPA modification. The source of post-translational modification by oxidation might be either enzymatic, such as tyrosinase in case of Tyr, or by reactive oxygen species (ROS), such as molecular oxygen (O_2_) or hydroxyl radicals (OH·). (**B**) The mean oxidation occupancy (Occ) was calculated using the formula: Occ (%) = ((Tyr_ox_Phe_2Ox_ abundance)/(Tyr_ox_Phe_2Ox_ abundance + Imm(Tyr or Phe) abundance))*100 from n=4 biological replicates. Each biological replicate was measured in 3 independent label-free LC/MS measurements (*23, 28*). Data are shown as mean ± SEM with each technical replicate shown in symbols for the respective tissue. (**C**) From the measured collagenous peptides, we determined the listed additional modification sides (alpha chain and protein ID as indicated). The number of the modification site refers to the absolute number in the expressed polypeptide. Marked with an asterisk (*) is the Tyr side previously identified (*16*). (**D**) The motif enrichment was calculated using dagLogo (*29*) using only collagenous peptides. The central F/Y modification site is centered at position 0. The y-axis is differential amino acid usage (DAU, %), which is a relative measurement for the occurrences of a motif (motif enrichment analysis). The coloring scheme is by chemical properties of the side chains. (**E**) Projection of the modification sites on a collagen type I 3D model generated in ColBuilder (*24*) and visualized in Ubuntu Pymol (*30*). Residues Phe865 (I), Phe1102 (I), Phe1187 (I), Phe1189 (I), Tyr1185 (I), and Tyr1114(II) (Phe/Tyr depicted in red/magenta) are close to crosslink sites (depicted blue).

The chosen analytical technique to determine the fraction of oxidized residues is based on Fourier Transform Isotopic Ratio MS (FT isoR MS) (*22, 23*). It allows to determine the occupancy of single oxidations of Tyr and/or double oxidations of Phe residues by immonium ion analysis in the polypeptides of these tissues. Such a post-translational modification on these two amino acids will produce a peak at m/z 152.0712 in MS/MS with high energy collision dissociation (HCD = 50 %). The mean oxidation occupancy (Occ) was calculated using the formula: Occ(%) = ((Tyr_ox_Phe_2Ox_ abundance)/(Tyr_ox_Phe_2Ox_ abundance + Imm(Tyr or Phe) abundance))*100.

The analysis showed that both kinds of residues are oxidized to DOPA across all investigated tissues (Figure 2B). Tyr residues are the more prevalent source with a mean oxidation occupancy of 13.1 ± 2.3% (mean ± SD) for the tail, 11.0 ± 1.5% for Achilles’ tendon tissue, and 3.8 ± 1.3% for meniscus tissue. The fraction was in a similar regime to the peptide count of 5% on the previously described single-site Tyr1114DOPA modification (*16*). For the same tissues, the occupancy for Phe was only 2.4 ± 0.8% or lower 1.3 ± 0.6%, and 0.8 ± 0.1%, respectively.

Nevertheless, given the lower conversion rate of Phe compared to Tyr in collagen I, Phe oxidation still contributes to the overall DOPA content. The conversion rate likely reflects that Phe is less prone to be oxidized by reactive oxygen species (*21*). Even though meniscus samples were overall less modified, their individual modification sites showed highest peptide counts (up to 100% of peptide modified to DOPA), that is, meniscus was more stably modified (Supplementary Fig. S3A).

We next searched for DOPA modification sites in collagenous proteins and could identify fourteen novel modification sites (Figure 2C). Of twelve Phe_2Ox_ sites, eight were found in collagen type I chains (out of 27 Phe in alpha (I/II)). Here, in addition to the previously identified Tyr1114DOPA, we uncovered one more Tyr_Ox_ (out of nine Tyr in alpha (I/II)) at position 1185 in alpha (I) chain. Other modification sites were found in fibrillar types II and XI, as well as associated type XII (Figure 2C). In the current setup, we did not identify more sites for oxidation on Tyr, which can be explained by low sequence coverage for collagenous proteins especially around the crosslinks (*16*). Based on a motif enrichment analysis, -G-F/Y-P/M/Q/S-is the favored motif in which the F/Y residue is doubly oxidized (Figure 2D, Supplementary Fig. S3B). Overall, in the canonical-GXY-sequence of collagen I (*13*), the X position is the one modified - both for Tyr and Phe residues (Figure 2C+D).

In the structure of collagen I (*24, 25*), both Tyr and Phe can be found in the close vicinity of crosslinks (*16*). Here, we found that six out of ten of the identified DOPA sites are in the vicinity to the crosslink sites: Phe865DOPA (alpha I), Phe1102DOPA (alpha I), Phe1187DOPA (alpha I), Phe1189DOPA (alpha I) and Phe794DOPA (alpha II), Tyr1185DOPA (alpha I) and Tyr1114DOPA (alpha II, described earlier (*16*)) (Figure 2E); they are predicted to be potential rupture sites (*16*). The close vicinity enables rapid and potentially direct transfer of the primary radical originating from rupture to the spatially close DOPA sites.

DOPA modifications of Phe and Tyr are known both as intentional modifications of collagen-like proteins in mussels (*14*) but also as an adverse site product of protein oxidation (*26, 27*). If the DOPA modifications we observe in collagen I originate from enzymatic or non-enzymatic (e.g., ROS-mediated, Figure 2A) oxidation remains to be investigated.

### L-DOPA is an efficient radical scavenger

Having identified DOPA both as stable anion radical in stressed collagen tissue and ubiquitous post-translational modification in different collagen type I tissues, we aimed to better understand its radical scavenging capacity. 2,2’-Azino-bis (3-ethylbenzothiazoline-6-sulfonic acid) (ABTS), which upon activation at neutral pH forms a stable N-centered radical with a local λ_max_ = 734 nm (*31*) (Figure 3A, Supplementary Fig. S4), was used to quantify the antioxidant characteristics of amino acids L-DOPA and L-Tyr. The two amino acids were tested in comparison to the known antioxidants ascorbate and Trolox. The latter was used in the past as the reference to compare the radical scavenging capacity of different substances (*31, 32*). Similar to L-DOPA, the hydroxyl groups in Trolox are oxidized yielding the respective quinone (*33*), while ascorbate is converted to its diketone dehydroascorbic acid (*34*).

**Figure 3.**
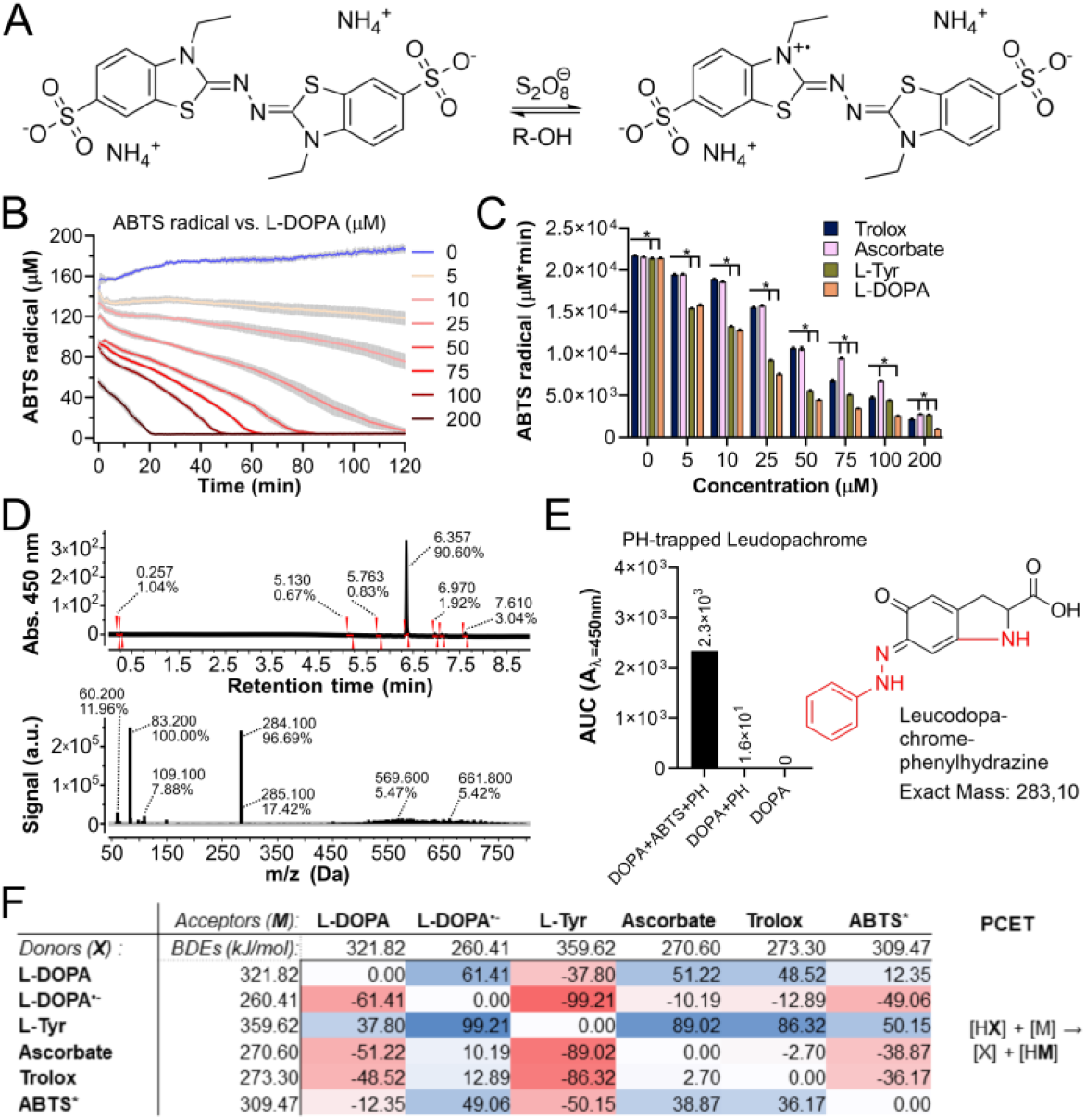
L-DOPA is an efficient radical scavenger. (**A**) The simplified reaction scheme shows the activation of 2,2’-Azino-bis (3-ethylbenzothiazoline-6-sulfonic acid) (ABTS) by oxidant ammonium persulfate (S_4_O_8_^2-^). The persulfate behaves in the reaction like a one-electron oxidant with a rate constant of 0.63 M^-1^s^-1^ (*36*). The ABTS radical is a strong chromophore with λmax=734 nm. The radical can then be quenched by an antioxidant (R-OH). (**B**) The reaction coordinate of the ABTS radical scavenging was measured as O.D. at λ=734 nm. The O.D. was converted to the ABTS (μM) by the conversion factor y=y*k (k=138.652). Recording in the plate reader started one minute after adding the reactants, because of the pipetting and the mechanics of the plate reader. The data represent n=4 independent measurements of each two technical replicates (mean ± SEM). (**C**) The ABTS radical concentration was determined as area under curve (AUC) analysis for the whole-time scale of each substance and concentration. For the given coordinates, AUCs were grouped according to the concentration (n=4*2 tech. repl., mean ± SEM). Significance (indicated by asterisk, p<0.05, difference from Trolox) was calculated by multiple t-tests (Holm-Sidak method, no consistent SD, α=0.05, for normality test and p-values, see Fig. S4). (**D**) The reaction product (phenylhydrazine (PH) adduct, exact mass 283.1 m/z, detected at with absorbance (Abs.) 450 nm at 6.357 min) was measured by HPLC/MS with ESI. The data was processed with MestReNova (v. 14.2.1) software (see also Methods section). (**E**) The oxidation product of L-DOPA after ABTS assay, circularized leucodopachrome (*35, 37*), was stabilized with phenylhydrazine and measured by LC-MS. Oxidation based changes as compared to L-DOPA were indicated in red. (**F**) In quantum mechanical calculation, the stability of radicals in the assay was calculated with DOPA anion radical (L-DOPA^·-^) having the lowest bond dissociation energy (BDE) of 260 kJ/mol. The values in the table indicate the differences in BDEs. The calculated mechanism of the proton-coupled electron transfer (PCET) is indicated on the right-hand side in the table. *BDEs of ABTS were calculated without an isodesmic correction.

The assay showed that L-DOPA scavenges ABTS radicals in a concentration dependent manner as does Trolox (Figure 3B+C). L-DOPA quenched the radical signal significantly faster as compared to Trolox and ascorbic acid at any given concentration stage (Supplementary Fig. S4B-C), as evident from the area under the curve of the ABTS radical concentration over time (Figure 3C). Notably, L-DOPA was also more efficient than L-Tyr in concentrations ≥10 μM in the current setup. All substances quenched ABTS radicals almost completely at the highest concentration levels. The first minute of kinetic information was not recorded due to the initial addition of ABTS and the mechanics of the plate reader.

L-DOPA quenched the radicals at a substoichiometric level: even at concentrations as low as 25 μM L-DOPA has almost completely consumed ~160 μM ABTS radicals within the given time frame (Figure 3B). The reason for that might be the complex chemistry of L-DOPA: it undergoes several conversion steps upon oxidation, including the formation of circularized products (*35*). In the first step of this mechanism, the primary amino group of L-DOPA attacks the phenol ring forming indolic leucodopachrome. Indeed, we could stabilize and detect this first product by addition of excess phenylhydrazine which specifically reacts with quinones (Figure 3D+E). This circularized product and others allow recovery of the hydroxyl groups for further oxidation steps, which is equivalent to a stronger antioxidant activity in the assay.

We quantified the radical stabilization energies of the different substances using quantum chemical calculations and confirmed that the DOPA radical anion is the most stable species (Figure 3F). This is in line with our EPR data and explains the absence of a tyrosine radical in stressed collagen. Our results also suggest DOPA-like molecules as highly potent antioxidants even outperforming ascorbic acid.

In the peptides, the primary amino group of L-DOPA is part of the peptide bond and not available for the circularization reaction. To better understand the behavior of peptide-bound DOPA, we synthesized a tripeptide based on the DOPA site originally identified in rat tail collagen (*16*): Glycine-X-aspartate (G-X-D), with X being either phenylalanine, tyrosine, or DOPA, which carry no, one or two hydroxyl groups, respectively (Figure 4A).

**Figure 4.**
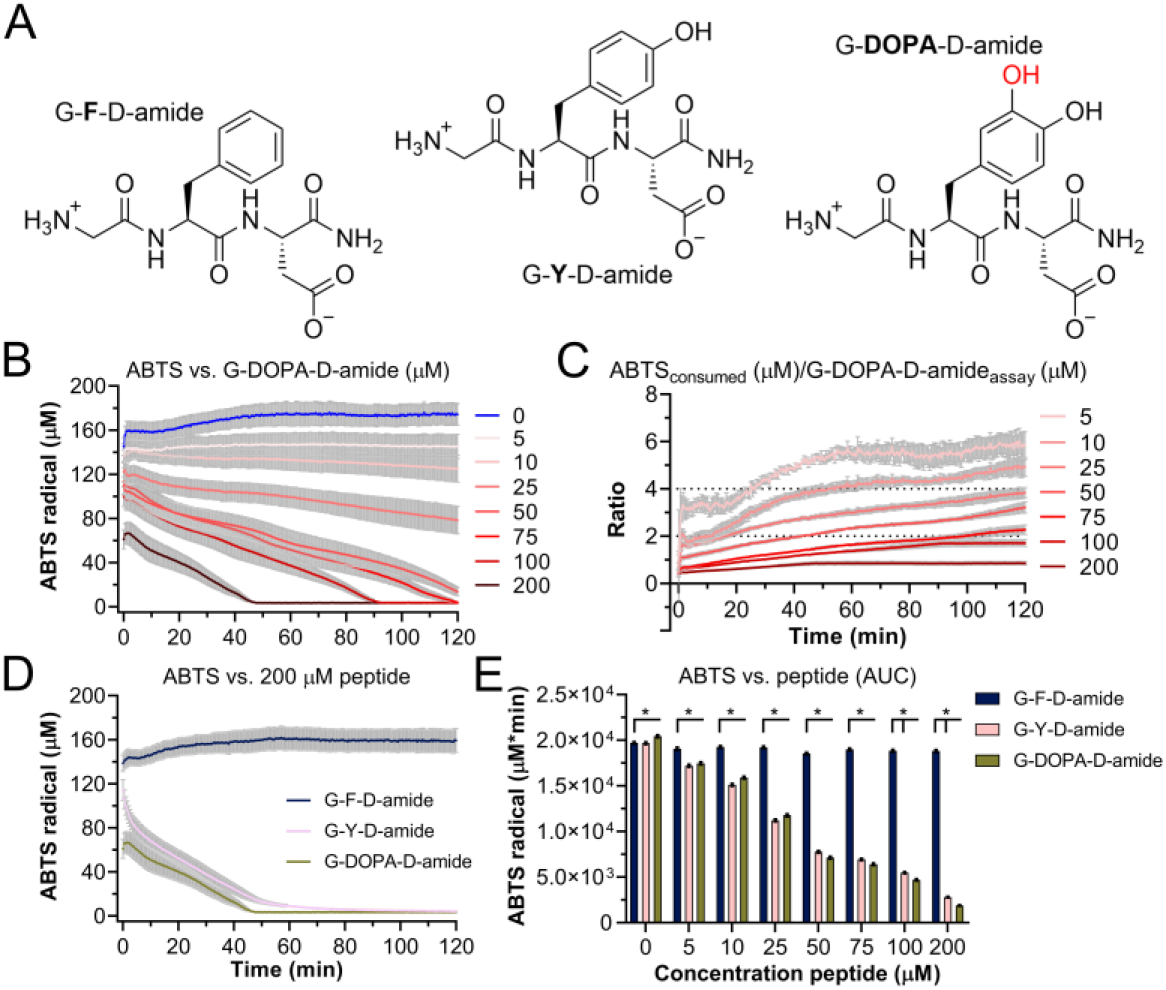
DOPA peptides scavenge radicals more efficiently than Tyr peptides. (**A**) Tripeptides were synthesized based on the earlier identified Tyr1029DOPA modification in collagen type I, alpha II chain (*16*). (**B**) ABTS radical concentration over time when scavenged by the G-DOPA-D-amide at different initial concentrations. The O.D. measured at λ=734 nm was converted to the ABTS radical concentration (μM) by the conversion factor y=y*k (k=138.652). The data represent n=4 independent measurements of each two technical replicates (mean ± SEM). (**C**) The ratio between consumed ABTS (calculated averaging the tech. replicates and subtracting them from the respective c=0 μM) and DOPA peptides was calculated by dividing the concentration of consumed radical by the assay concentration of the scavenger. (**D**) The reaction coordinate of the ABTS radical scavenging assay at half-maximum concentration of the peptides. The O.D. 734 nm was converted to the ABTS (μM) by the conversion factor y=y*k (k=138.652). The data represent n=4 independent measurements of each two technical replicates (mean ± SEM). (**E**) The ABTS radical concentration was determined as area under curve (AUC) analysis for the whole-time scale of each substance and concentration. For the given coordinates, AUCs were grouped according to the concentration (n=4*2 tech. repl., mean ± SEM). Significance (indicated by asterisk, p<0.05, difference from G-DOPA-D-amide) was calculated by multiple t-tests (Holm-Sidak method, no consistent SD, α=0.05, for normality test and p-values, see Fig. S5).

As for the single amino acid, the Gly-DOPA-Asp-amide peptide (G-DOPA-D-amide) scavenged the ABTS radicals in a concentration dependent manner (Figure 4B). Both the DOPA and Tyr containing peptides (G-Y-D-amide) showed radical scavenging capacity, in sharp contrast to Gly-Phe-Asp-amide (G-F-D-amide) (Figure 4D+E, Supplementary Fig. S5). When comparing the two active peptides both in their kinetics and total amount of radical degradation (shown as AUC), the DOPA peptide was the more efficient in quenching ABTS at higher concentrations (≥100 μM, multiple t-test, p<0.05). We also observed the previously described adduct of L-Tyr with an ABTS radical (*38*) which, however, was absent in the L-DOPA reaction (Supplementary Fig. S6). Phe residues did not show any activity and thus do not contribute to the radical scavenging in the collagen polypeptide - unless they are oxidized (Figure 2).

Our joint EPR, MS and radical scavenger kinetic data suggest that Phe, Tyr and DOPA might work together as a collective buffer system in collagen: Phe or Tyr residues, as the primary source of DOPA modifications, can be oxidized in an initial scavenging event. DOPA as a newly formed post-translational modification works itself as a radical scavenger in collagen and remains a fingerprint of mechano-oxidative stress within the polypeptide. Hence, we propose DOPA to be not only as an adverse side product of oxidation but also a beneficial radical scavenger in the collagen I tissue.

The ratio between the added peptides and the ABTS radicals they scavenged was mostly exceeding 1:2 (except for measurements with excess of peptide), the ratio expected after complete oxidation of DOPA by two electrons to its quinone. (Figure 4C). This substoichiometry is in line with the radical scavenging capacity already observed for the amino acid DOPA (Figure 3B), which we attribute to a temporary recovery of DOPA from its quinone due to slower secondary reactions with ABTS.

### DOPA radical scavenging produced peroxide

DOPA is a very efficient radical scavenger as it is a stable radical itself. However, most organic radicals can only be temporarily stabilized before they are converted further. In rat tail tendons, mechanical stress led to increased hydrogen peroxide concentrations (*16*), suggesting that DOPA radicals in collagen are converted to peroxide in the presence of molecular oxygen/superoxide. To put this scenario of radical degradation via DOPA to test, we followed the formation of hydrogen peroxide after the reaction of ABTS radical scavenging by the DOPA tripeptide. We utilized a boric acid-based chemiluminescence probe (Lumigen HyPerBlu™) to detect hydrogen peroxide specifically and directly in the assay without further cleanup (Figure 5A). We took samples directly from the ABTS assay after the reaction was complete and again compared G-X-D-amide tripeptides containing Phe, Tyr or DOPA.

**Figure 5.**
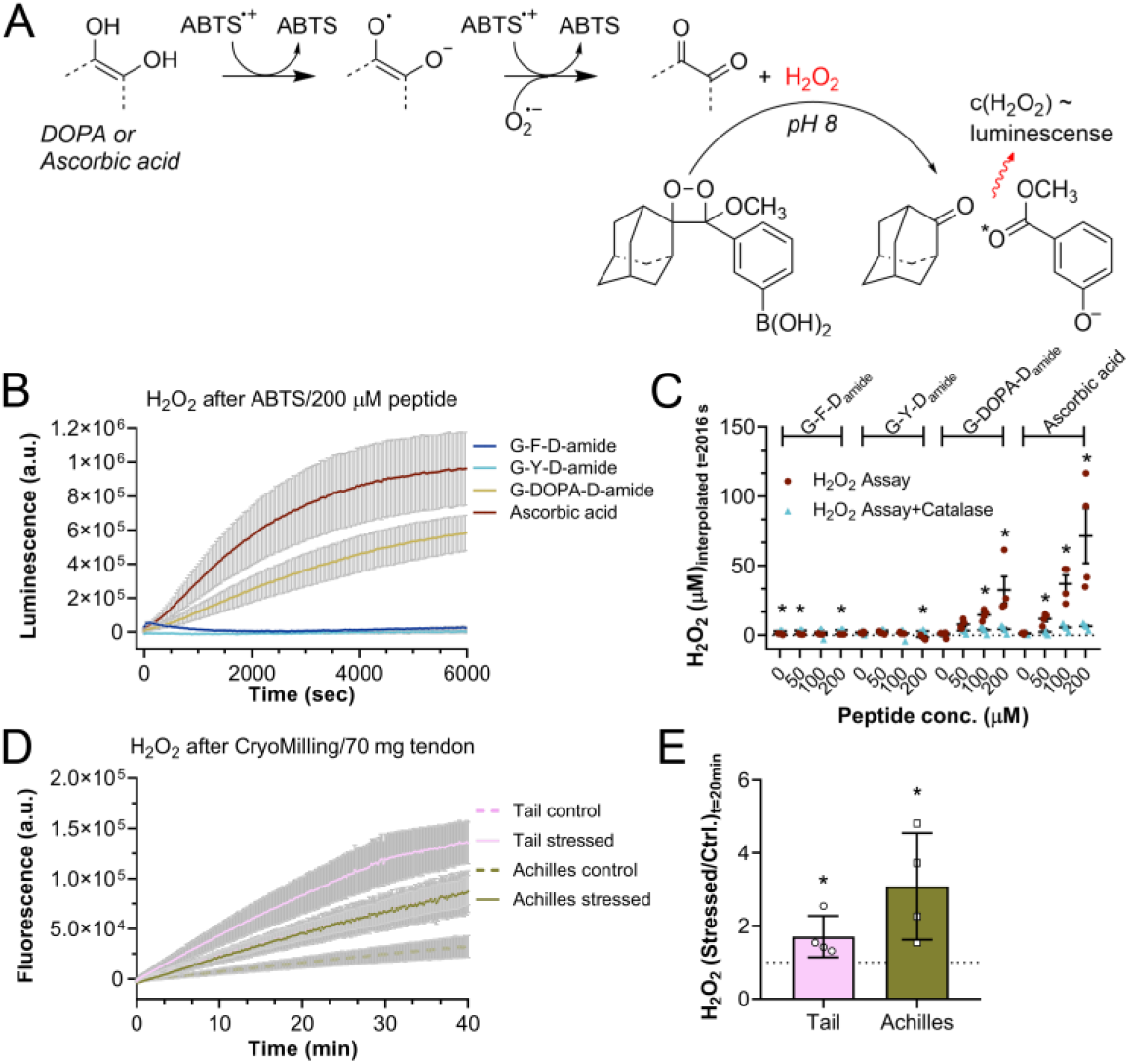
Hydrogen peroxide is the end product of the DOPA pathway. (**A**) Simplified reaction scheme for the formation of hydrogen peroxide (H_2_O_2_) after quenching the ABTS radical (ABTS^·+^) by ascorbic acid or DOPA. The boric acid-based luminescence probe HyPerBlu^™^ reacts with H_2_O_2_ under formation of a chemiluminescence signal. (**B**) H_2_O_2_ detection by HyPerBlu^™^ after ABTS assay with ascorbic acid or peptides at 200 μM. Samples were taken directly from ABTS assays and luminescence recorded for 6000 s at intervals of 42 s (n=4 experiments with each 2 tech. replicates, mean ± SEM). (**C**) H_2_O_2_ concentrations estimated at t=2016 s (approx. peak point of detection for 200 μM H_2_O_2_ standard) by interpolation of the respective standard curve with background (c_H2O2_=0 μM) corrected values. The data represent n = 4 independent assays of each 2 tech. replicates (mean ± SEM). Significance between assay and assay with catalase is indicated with an asterisk (*) and was tested by non-parametric Mann-Whitney test (one-sided, 95% confidence level). (**D**) In tissue, H_2_O_2_ was detected by horseradish peroxidase assay using a near-infrared dye. The kinetic was recorded by fluorescence signal (Ex: 640 nm, Em: 680 nm) in untreated samples (“Control”) and cryomilled (“Stressed”) samples (both ~70 mg) for 40 min (n=4 with each 2 tech. replicates, mean ± SEM). Signals were background corrected by deducing fluorescence signals in an empty tube. (**E**) The ratio between stressed and unstressed samples was calculated at t=20 min by dividing the background corrected signals. For tail and Achilles’ tendons, the values diverge significantly from 1 (Shapiro-Wilk test for normality, p=0.0613 & 0.7867, respectively; unpaired t-test with 95% confidence level, one-sided, Welch’s correction, p=0.0441 & 0.0325, respectively).

Among the different peptides, only the DOPA peptide showed an ABTS concentration dependent increase in the luminescence signal over time (Fig. 5B-C, Suppl Fig. S7–9). Remarkably, hydrogen peroxide production through DOPA radical scavenging nearly reaches levels observed for ascorbic acid, which is well known to generate hydrogen peroxide during radical scavenging (*39*) (Fig. 5A). In contrast, the G-Y-D-amide scavenged ABTS radicals (compare Figure 4D) but did not show any concentration dependent peroxide production (Figure 5B-C and Suppl Fig. S7–9). The Phe tripeptide showed activity in either assay only at background level. The HyPerBlu chemiluminescence signal is very specific for hydrogen peroxide, as demonstrated by adding catalase, which fully quenched the luminescence signal over the whole assay time of 6000 s (Figure 5C and Suppl Fig. S7). For higher concentrations of DOPA peptide as well as ascorbic acid, the peroxide concentration in the assay was significantly elevated over the catalase control (Figure 5C, Supplementary table 2). For the Phe peptide, the peroxide concentration of the catalase control was significantly elevated but not in a concentration dependent manner.

The hydrogen peroxide measurement demonstrated that DOPA peptides shared the mechanism of radical scavenging with ascorbic acid, underlining that the modification might be beneficial for mechanoradical scavenging in fibers. Notably, the ratio between hydrogen peroxide and DOPA peptide after ABTS assay was ~1:6, with this value being approximately constant for 50 μM, 100 μM & 200 μM DOPA peptide to 7.8 μM, 14.6 μM & 32.6 μM peroxide, respectively (Figure 5C and Suppl Fig. S9). As each DOPA can quench two radicals and convert those to one hydrogen peroxide, we expected a 1:1 ratio instead. Thus, yet unknown competitive mechanisms might play a role here, possibly due to the complex chemistry of ABTS (*40, 41*).

In the next step, we went back to tendon samples, in order to test if the previous observation of hydrogen peroxide production after mechanical stress in tail tendons extended also to other collagen tissues (*16*). For the stress condition, we used cryomilling to exclude secondary effects like denaturation of the fibers and to ensure sufficient signal during the measurement. In the assay, we measured hydrogen peroxide in ground tissue in comparison to untreated samples using a fluorescence horseradish peroxidase assay (Figure 5D, Suppl Fig. S10). A significant increase in fluorescence was observed for tail and Achilles tendon upon mechanical treatment; due to the low amounts no successful detection was possible with meniscus (Figure 5E, Suppl Fig. S10E-H). Overall, we conclude that the formation of hydrogen peroxide is a general consequence of mechanical stress in collagen tissues and can be attributed to the presence of DOPA as a post-translational modification of the collagen polypeptide (Figure 2B).

### DOPA is oxidized to its quinone but not to Topa

Hydrogen peroxide formation as a product of the DOPA pathway should also involve the full oxidation of DOPA to its quinone. While the quinone itself proved difficult to measure directly in peptides by mass spectrometry, we stabilized it with phenylhydrazone which forms a stable adduct (*37, 42*) (Figure 6A). The adduct was detected at an exact mass of 456 Da corresponding to the DOPA quinone peptide plus phenylhydrazone (−2 H) (Figure 6B). We were not able to detect triple oxidized phenylalanine residues, trihydroxyphenylalanine (Topa), in our DOPA peptides, in contrast to previous studies using the same labeling technique (*39*). The DOPA quinone adduct was only found in ABTS oxidized peptides while in its absence, labeling was almost at the background level (Figure 6C). Thus, DOPA oxidation to its quinone is a consequence of the formation of the DOPA radical, i.e., its semi-quinone.

**Figure 6.**
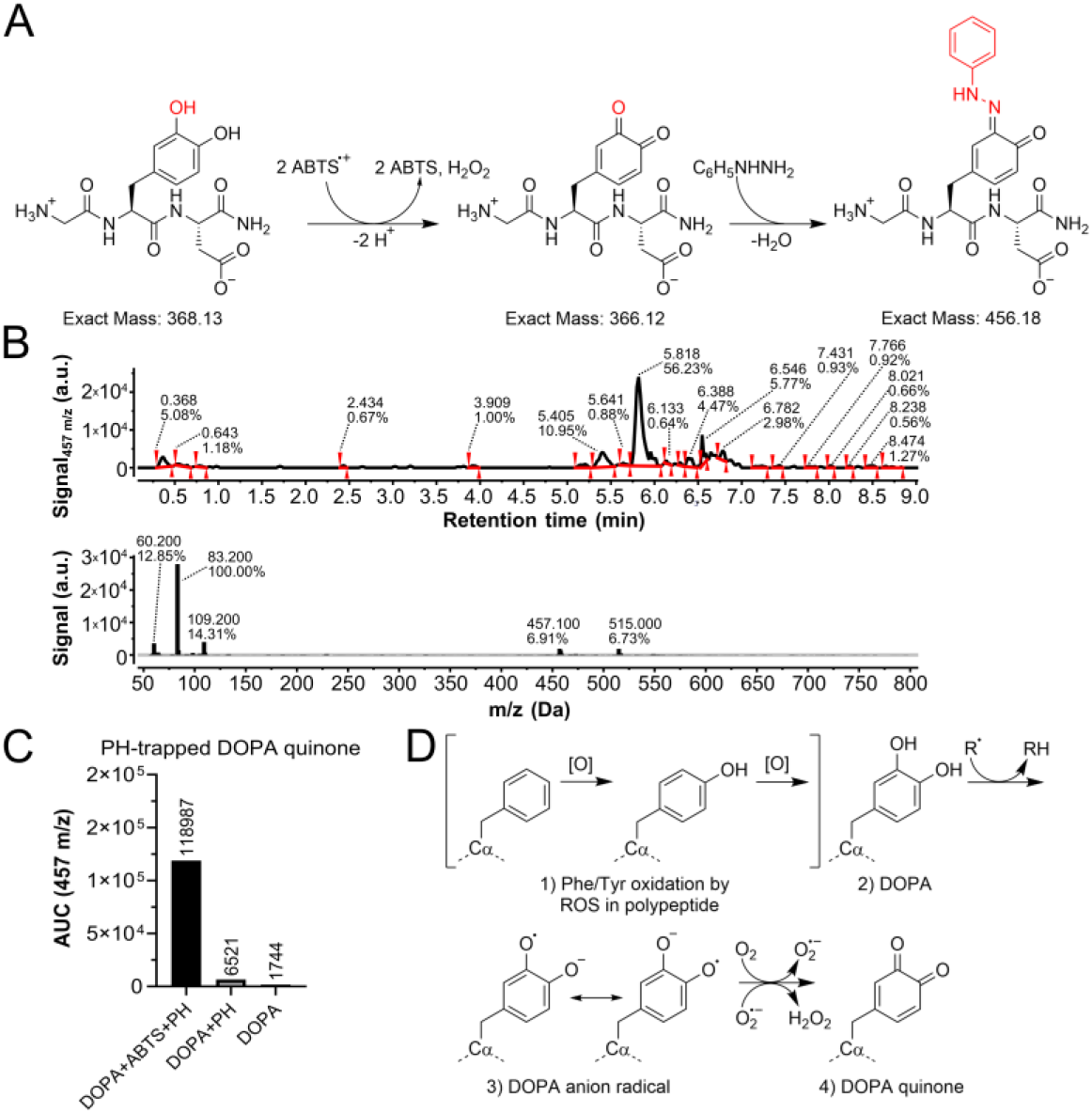
DOPA is oxidized to its quinone during radical scavenging. (**A**) Reaction scheme for DOPA oxidation in ABTS assay with subsequent phenylhydrazine (PH) (C6H5NHNH2) modification of the quinone moiety. The stabilized quinone has an exact mass of ~456 Da. (**B**) After ABTS assay with 500 μM radical and 500 μM peptide, a peptide-phenylhydrazine adduct was detected at the predicted mass of 456 Da. The reaction products were measured by HPLC/MS with ESI. The data was processed with MestReNova (v. 14.2.1) software (see also Methods section). (**C**) Comparing HZ-modified DOPA peptide samples from ABTS assay (DOPA+ABTS+HZ) with mock incubated samples (DOPA+HZ) and unmodified DOPA peptides (DOPA), we found x-fold increase of the adduct as compared to the controls as measured by the area under curve (AUC). (**D**) Initially, the peptide-bound Phe or Tyr residues are oxidized [O] to DOPA by reactive oxygen species (ROS). Radicals (R), either mechano- or ABTS-radicals, are then scavenged by DOPA, which is finally oxidized to its quinone and hydrogen peroxide (H_2_O_2_). DOPA (quinone) remains as a molecular fingerprint in the collagen (poly-)peptide.

Our data based on the ABTS assay thus support the mechanism previously hypothesized for the redox reactions in collagen fibers (*16*): Peptide-bound Phe or Tyr is first oxidized to DOPA (compare Fig. 2+6D), and DOPA is then further oxidized to its quinone. Here, DOPA scavenges both mechano- and ABTS radicals in a similar fashion (Figure 6D). In total, the Tyr residue can undergo three oxidation events or donate three electrons. A key product of this radical scavenging is hydrogen peroxide while DOPA (or its quinone) remains a molecular fingerprint in the collagen polypeptide (*27*).

In the assay, the source of substoichiometric consumption of ABTS radicals by DOPA are probably secondary reactions of ABTS or its degradation products. As mentioned above, we do not find evidence for the oxidation of DOPA to Topa as a reason. In situ, the DOPA quinone might be recovered for instance by GSH in a mechanism similar to the one of ascorbic acid (*34*).

## Conclusion

By combining EPR spectroscopy and mass spectrometry with ex-situ tissue measurements and biochemical assays, we confirmed and expanded our hypothesis on DOPA’s role in collagen type I tissue. Here, we present evidence that DOPA is a stable modification of collagen tissue which is also found in the physiologically relevant tissue of the Achilles’ tendon and cartilage from meniscus. In these tissues, Tyr residues are the primary source of DOPA though also Phe appeared as a minor source. DOPA and likely also its precursor Tyr are part of a buffer system in collagen that scavenges radicals that are for instance generated during homolytic bond rupture during mechanical stress. Tyr residues are then in the first step oxidized to DOPA which subsequently acts itself also as a radical scavenger in the tissue. The products of radical scavenging by DOPA are hydrogen peroxide, in direct analogy to and with an efficiency approaching that of ascorbic acid as a radical scavenger, and DOPA quinone. While we find Tyr to have a radical scavenging capacity similar to that of DOPA, it can only marginally contribute to hydrogen peroxide formation. Our data points towards a Janus-faced role of DOPA which is both a product of oxidative stress and at the same time helps to stop further radical chain propagation, either by stabilizing free radicals as DOPA radicals or by converting radicals to the molecular product H_2_O_2_, which could be removed by various cellular systems. DOPA has been known as post translational modification with a role as an adhesive in collagen-like proteins of mussels for a long time (*14*). In mussel bypass threads as well as in the present collagen samples, the source of DOPA is unclear. There are three potential ways to incorporate the modification: 1) modification of Tyr (or Phe) by oxidation **(*21, 27*)**, 2) enzymatic conversion of Tyr by the enzyme tyrosinase (*18*), and 3) during protein synthesis by tRNAs usually carrying Phe or Tyr **(*43*)**. We assume that the first path is the primary source of DOPA in collagen tissue as it would fit to a role of Tyr and DOPA as a buffer system for mechanical and hence oxidative stress. DOPA then can serve as a fingerprint for stress and age. Interestingly, for Achilles’ tissue, it has been noted early on that Tyr is lost with age **(*44*)**. In collagen polypeptides, DOPA might also help to assess the state of a tissue in health and disease. The other question arising from our study is the rate of DOPA turnover, i.e., if DOPA quinone can be reduced to DOPA to allow for another round of redox cycling.

Taken together, our study discovers DOPA as a component of tendons and meniscus and potentially many other connective tissues and identifies these unique protein building blocks as an efficient antioxidant and redox relay system that converts radicals into hydrogen peroxide. We propose that this novel defense mechanism built into connective tissue can guide the design of biocompatible materials and serve as a therapeutic target in oxidative stress related diseases and aging.

## Materials and methods

### X-band EPR spectroscopy

The X-band EPR spectroscopy was carried out as previously described (*16*). Collagen tissue from Rattus norvegicus was a donation of the IBF unit at Heidelberg University (Germany). Tendons from the tail and Achilles’ were cut into pieces of 2-3 cm in length, 2-3 mm^2^ in diameter with weights between 15-25 mg. Meniscus tissue was cut from the knee. Achilles and Meniscus samples were briefly washed in demineralized water to remove major contaminations. All tissue samples were equilibrated at room temperature before the experiment. The tissue was then measured in 4 mm (O.D.) quartz tubes (Wilmad-LabGlass, USA, 707-SQ-x) once before and once after pulling at X-band (~9 GHz microwave frequency) with a Bruker EPR ESP300 or EMXmicro setup using standard parameters (2 mW microwave power, 0.2 mT modulation amplitude, 100 kHz modulation frequency, 20 scans/sample with 60 s sweep time). The pulling experiment was conducted on a LEX810 (Dia-Stron, USA) with stress-strain curve recorded by UvWin v. 3.0 software. Tendons were pulled at constant force of 15 gmf for 1000 s. The creep rate was set to 0.01 mm/s. Both EPR spectra and strain measurements were analyzed using R statistics (*45–47*).

### G-band EPR spectroscopy

All biological samples were prepared as described for the X-band experiments. Pulsed EPR measurements at 180 GHz frequency were performed at 40 K on a home-built G-band EPR spectrometer (*48*) with a microwave power of about 100 mW. For the experiments, tendon collagen samples crushed at room temperature were transferred into quartz tubes (I.D. 0.4 mm, O.D. 0.55 mm). The field-swept Hahn echo-detected EPR spectra were recorded with pulse lengths of 42 ns and 72 ns. The pulse separation time was set to 200 ns. Shot repetition time was 10 ms and the number of shots per point was 1000.

The experimental X-band and G-band EPR spectra were simulated using Easyspin software (*20*) in the solid state.

### Mass spectrometry of collagen tissue

Collagen tissue from Rattus norvegicus (8 weeks old, female) was a donation of the IBF unit at Heidelberg University (Germany). Tissue samples (Achilles tendons, tail tendons, meniscus) were collected into ice-cold Dulbecco’s phosphate buffered saline (Sigma, D8537). They were then briefly washed with deionized water and defatted with 100% ethanol and 70% isopropanol. Dried samples were then dissolved to a concentration of 1 mg/ml in 0.02 N acetic acid for 72 h under agitation; after incubation, the solution was lyophilized.

Freeze-dried collagen samples were dissolved in 50 mM ammonium bicarbonate (approximately 200 μg per each) at 70 °C for 3 h. Later samples were centrifuged at 14.000 x g and the supernatant was taken as the solubilized collagen. The amount of collagen in each sample was assessed by Pierce™ BCA Protein Assay Kit. Collagens were then digested using Sequencing Grade Modified Trypsin from Promega with a ratio of 1:50 (trypsin: collagen) overnight at 37 °C. Samples were acidified with formic acid (98%) in order to quench the digestion process. Samples desalting was carried out using HyperSep Filter Plates, C18 (Thermo Scientific).

Peptides were reconstituted in solvent A (98/1.9/0.1% water/acetonitrile/formic acid) and approximately, one μg samples injected on a 50 cm long EASY-Spray C18 column (Thermo Fisher Scientific) connected to an Ultimate 3000 nanoUPLC system (Thermo Fisher Scientific) using a 75 min long gradient: 4-15% of solvent B (98% acetonitrile, 0.1% FA) in 50 min, 15-35% in 10 min, 35-95% in 3 min, 95% for 7 min, 95-4% in 1 min, and 4% of solvent B for 4 min at a flow rate of 300 nL/min. The targeted tandem mass spectra (tMS4) were acquired on Orbitrap Fusion Lumos Tribrid mass spectrometer (Thermo Fisher Scientific) ranging from m/z 50 to 200 at a resolution of R=60,000 (at m/z 200) targeting 5×104 ions for maximum injection time of 10 ms, with isolating precursor ion at 800 m/z with charge state of 2+ and isolation width of 1000 Th in the quadrupole. The higher-energy collisional dissociation (HCD) was set at 50%.

Using a technique similar to Fourier Transform Isotopic Ratio Mass Spectrometry (FT isoR MS) (*22, 23*), we determined the occupancy of single oxidation modification on Tyr and/or double oxidation of Phe residues by immonium ion analysis. Such a modification on these two amino acids will produce in MS/MS with high energy collision dissociation (HCD = 50%) a peak (-> Y_ox_F_ox2_) at m/z 152.0712. Oxidation Occupancy (Occ) was calculated using the formula: Occupancy (%) = ((Y_ox_F_ox2_ abundance)/(Y_ox_F_ox2_ abundance + Imm(Y or F) abundance))*100.

### ABTS radical scavenging assay

The radical scavenger assay using 2,2’-Azino-bis(3-ethylbenzothiazoline-6-sulfonic acid) diammonium salt (ABTS) was conducted as described previously (*31*) with modifications. In brief ABTS tablets (Sigma-Aldrich, A9941) were dissolved in Dulbeccos phosphate buffered saline (PBS) (Gibco, 14190-094) to a final concentration of 10 mM. ABTS was then activated using ammonium persulfate (Carl Roth, 9592.3) with a final concentration of 7 mM and 2.45 mM, respectively. The reaction was incubated in the dark for 1 h. Small molecules (L-DOPA (Sigma-Aldrich, D9628), L-Tyr (Sigma-Aldrich, T8566), L-Phe (Sigma Aldrich, 78019), Trolox (Sigma Aldrich, 238813), Ascorbate (Sigma Aldrich, 255564)) were freshly dissolved to a concentration of 1 mM in PBS. Likewise, custom-synthesized tripeptides (G-Y-D-amide, G-DOPA-D-amide, G-F-D-amide, cleaned-up as acetate, based on sequence described earlier (*16*)) by PSL GmbH (Heidelberg) were dissolved to a concentration of 1 mM, aliquoted for one-time use and stored at −80 °C. For the different concentrations, the stocks were diluted to 5x assay concentration. The kinetic measurement was conducted in PBS with a total volume of 200 μl (96-well flat bottom plate, transparent) using assay concentrations of 2.8 mM ABTS radical and small molecules as indicated. Absorbance as O.D. at λ=734 nm was measured for 120 min with one measurement point every 30 s (BMG, Fluostar Omega).

### Hydrogen peroxide assays

The hydrogen peroxide was quantified using HyPerBlu (Lumigen, A97286) content after ABTS radical scavenging assay (see also Methods “ABTS radical scavenging assay”). The ABTS radical scavenger assay was conducted as described using assay concentrations of 0 μM, 50 μM, 100 μM and 200 μM peptide or ascorbate in duplicate. After 2 h, an aliquot of 10 μl from each well was then transferred into a white 96-well flat bottom plate with 10x diluted HyPerBlu in 200 mM NaH2PO4/NaH2PO4 buffer, pH 8. The hydrogen peroxide standard curve was pipetted on the same plate from 1 mM working stock prepared on the day of the measurement. Chemiluminescence of HyPerBlu reaction with hydrogen peroxide was recorded for 100 min on (BMG, Pherastar) with one measurement point every 42 s. The luminescence at the indicated time point was used to interpolate concentrations in assays from the peroxide standard curve.

In tissues, hydrogen peroxide was quantified by a fluorometric near infrared horseradish peroxidase kit (Abcam ab138886). The detection was carried out in all-black, F-/solid bottom, low-binding 96-well plates (Greiner bio-one) using the top-read function on CLARIOstar 430-0671 (BMG). All assay components were diluted according to manufacturers’ instructions and frozen at −80 °C until further use, except for the hydrogen peroxide standard which was stored at 4 °C. The hydrogen peroxide standard was diluted in kits’ assay buffer. Collagen tissues were prepared from animals between 3-6 months of age. ~70 mg sample for cyromilling were cut to pieces of 2-3 mm and grounded with zirconium dioxide marbles (ø=5 mm) in 2 ml reaction tube on Retsch Cyromill (4x 2:30 min) at −196 °C. Control samples were likewise transferred into a 2 ml reaction tube with two marbles but left untreated. Samples were then overlaid with 200 μl assay buffer and incubated under shaking for 5 min at room temperature. Samples and standards were then incubated 1:1 with the reaction mix. Fluorescence was recorded for 40 min every 10 s using λex=640 ± 10 nm, dichroic λ=660 nm, and λem=680 ± 15 nm.

### Mass spectrometry of phenyl hydrazine adducts

For the mass spectrometry measurement of peptides, quinones in peptides were stabilized using phenylhydrazine as described previously (*37*). The peptides were incubated with ABTS radical as described for the ABTS radical scavenging assay. Here, the peptides were diluted to an assay concentration of either 62.5 μM or 500 μM and exposed to 500 μM activated ABTS or PBS mock control. After 2 h of incubation with ABTS radical, phenylhydrazone was added to a final concentration of 5 mM.

Samples were then measured by HPLC/MS with ESI, Agilent 1260 Infinity system equipped with a 6120 Quadrupole mass detector and evaporative light scattering detector (ELSD). Separation was performed on Acidic Method Kinetex 2.6 μm C18 100 Å, LC column 50 x 2.1 mm; at 40 °C; with flow rate - 0.6 mL/min. Solvent “A” was 0.01% HCOOH in water. Solvent “B” was 0.01% HCOOH in MeCN. The method was: 100% A for 2 min, then from 100% to 10% A in 10 min, then 1% A for another 10-12 min. Data was processed with MestReNova (v. 14.2.1) software.

### Quantum mechanical calculations

BDEs were calculated using the hybrid functional BMK (*49*) with the 6-31+G(2df,p) basis set using the isodesmic reaction method. As experimental reference structure Phenol was used for Trolox and Ascorbate, for ABTS no experimental BDE was available, thus the BDE was computed from DFT directly. All experimental BDEs and reference structures are reported by Treyde et al. (submitted (*50*)). Quantum mechanical calculations were performed using Gaussian09 (*51*).

## Acknowledgements

We thank Dr. Aubry K. Miller for help with LC-MS measurements. We further thank Marina Bennati for helpful discussion on EPR data, and Christina Boroffka of the IBF Heidelberg for help with collagen tissue sampling.

We acknowledge funding through Klaus Tschira Foundation, the European Research Council (ERC) under the European Union’s Horizon 2020 research and innovation programme (grant agreement No. 101002812 to F.G., and 742039 to T.P.D.), the Deutsche Forschungsgemeinschaft (DFG, German Research Foundation) under Germany’s Excellence Strategy – 2082/1 – 390761711, the Federal Ministry of Education and Research (BMBF) and the Ministry of Science Baden-Württemberg within the framework of the Excellence Strategy of the Federal and State Governments of Germany, the state of Baden-Württemberg through bwHPC, as well as the DFG through grant INST 35/1134-1 FUGG.

## Supplementary information

### DOPA radicals form in different collagen type I tissues

**Figure S1.**
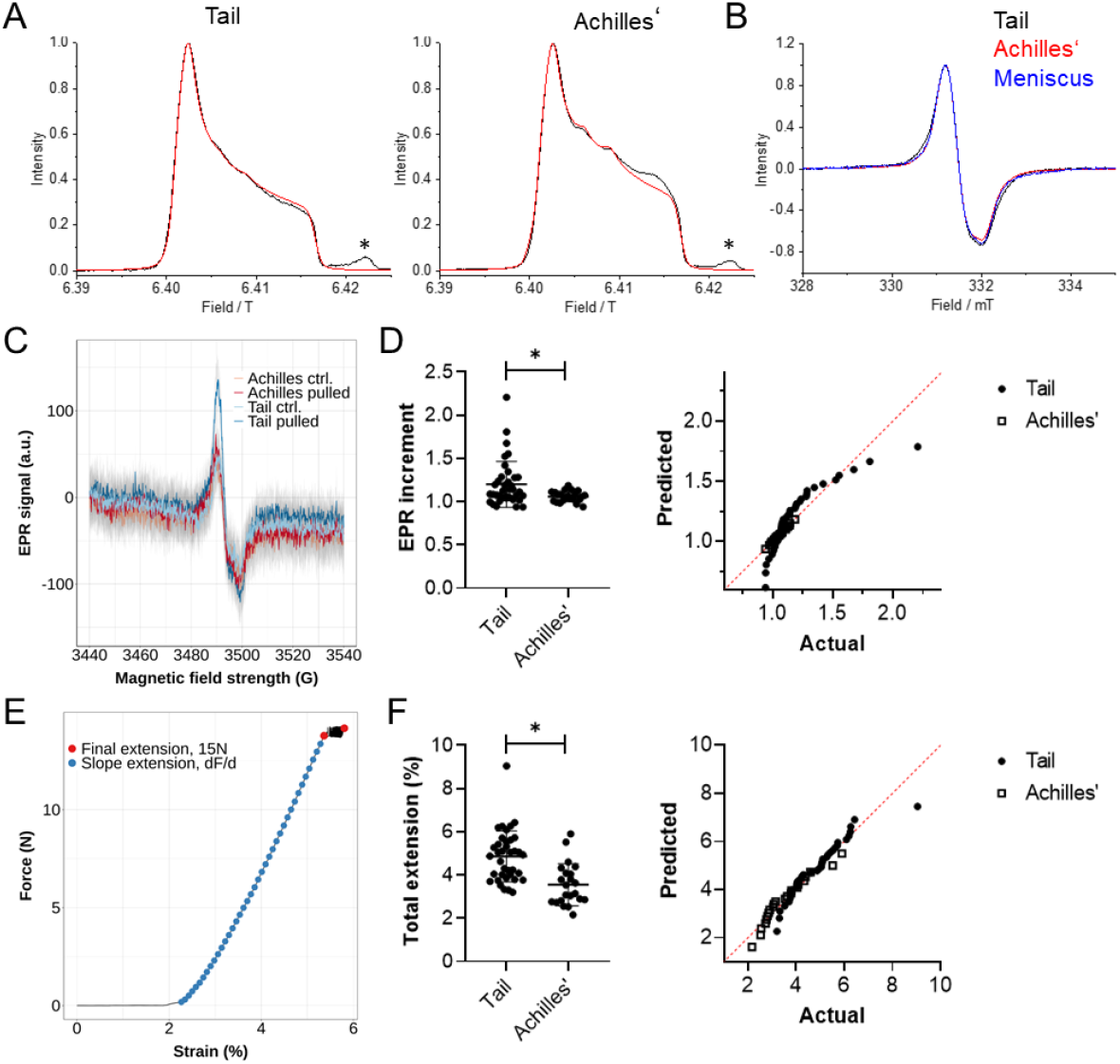
Strain and EPR measurements of collagen tissues. (**A**) G-band EPR spectra measured at 40 K and (**B**) X-band EPR spectra measured at 298 K for ground collagen tissue (Tail, Achilles’, samples 2, black curves) showed a similar radical signature (each curve represents one biological replicate, n=2). Fitting with EasySpin (*20*) (simulation shown as red curves) showed DOPA anion radical (g_x_=2.0071, g_y_=2.0066, g_z_=2.0022, 85% and 65% for tail and Achilles’, respectively) as well as two unknown organic radicals (g_x_=2.0075, g_y_=2.0055, g_z_=2.0021 and g_x_=2.0075, g_y_=2.0046, g_z_=2.0021). The signal marked by * in the G-band spectra in (A) appears from the quartz capillary. (**C**) The EPR signal for each tendon sample was measured before (control/ctrl.) and after pulling (pulled) at constant force. Shown here are representative curves given as mean ± standard error from n=20 scans of the given tissue sample (**D**) For the EPR signal increment, first the difference between minimum and maximum of the signal depicted in (A) was determined. From the differences, or peak sizes, before and after pulling the EPR signal increment was calculated as quotient between these values. Here, we report the values for the tail tendons (n=37 measurements) with 1.20 ± 0.27 (mean ± standard deviation), for the Achilles’ tendons (n=22 measurements) with 1.06 ± 0.06. In the Shapiro-Wilk test, the tail data set was not normally distributed (p<0,0001), while Achilles’ showed normal distribution (p=0.9985). The difference between the two tissues was significant in Mann-Whitney non-parametric test (two-tailed, p=0.0227). (**E**) For each pulling experiment, the stress strain curve was measured. Shown here is a representative curve for a constant force experiment. From the strain, the total extension was calculated from slope Δ>0.5 plus the final extension at 15 gmf. (**F**) On average, the extension was 4.86 ± 1.17% for tail, and 3.56 ± 0.97% for Achilles’ tendons. In the Shapiro-Wilk test, the tail data set was not normally distributed (p=0.0060), while Achilles’ showed normal distribution (p=0.1076). The difference between the two tissues was significant in Mann-Whitney non-parametric test (two-tailed, p<0,0001).

**Table S1.** EPR parameters for the DOPA from X-band (X) and G-band (G) measurements

| Sample | Contribution of DOPA radicals<br>($g_x=2.0071$; $g_y=2.0066$, $g_z=2.0022$) | Contribution of radical 2<br>($g_x=2.0075$; $g_y=2.0055$, $g_z=2.0021$) | Contribution of radical 3<br>($g_x=2.0075$; $g_y=2.0046$, $g_z=2.0021$) |
| --- | --- | --- | --- |
| Tail – sample 1 | 66.6 % | 16.6 % | 16.6 % |
| Tail – sample 2 | 84.6 % | 7.7 % | 7.7 % |
| Achilles – sample 1 | 57.1 % | 24.3 % | 18.6 % |
| Achilles – sample 2 | 64.5 % | 17.7 % | 17.7 % |
| Meniscus | 54.0 % | 23.0 % | 23.0 % |

**Figure S2.**
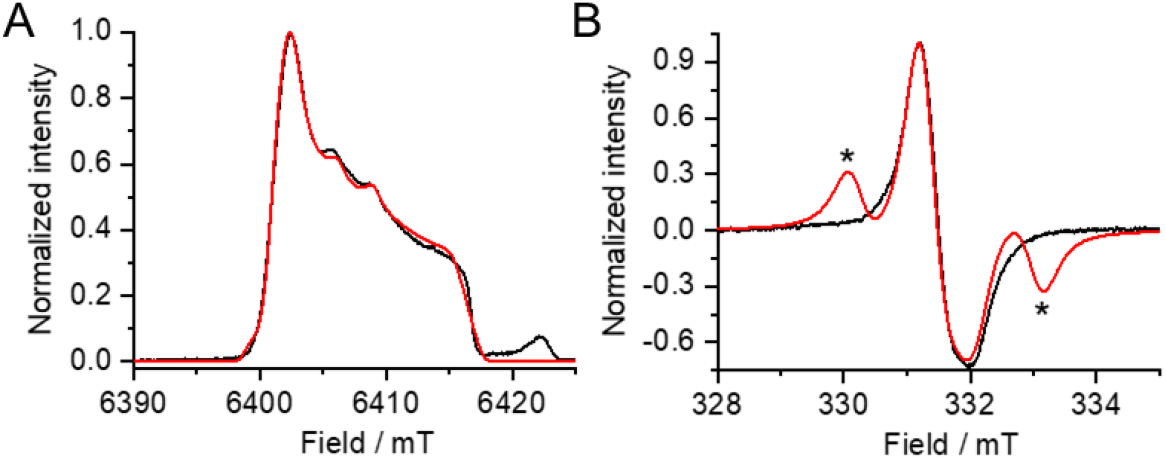
Simulation of a tyrosyl radical in the measured EPR spectra. (**A**) G-band EPR spectrum measured at 40 K for Tail collagen sample (black line). (**B**) X-band EPR spectrum (black line) measured at 298 K for the same collagen tissue mentioned in (A). Fitting was done with EasySpin (*20*) (simulation shown as red curves) assuming DOPA anion radical (g_x_=2.0071, g_y_=2.0066, g_z_=2.0022, 66.6%) and tyrosyl radical (g_x_=2.0074, g_y_=2.0050, g_z_=2.0022, A_x_=58 MHz, A_y_=80 MHz, A_z_=52 MHz, 33.3%). This assumption allows quite well getting experimental shape of G-band EPR spectrum, however X-band CW EPR spectrum does not demonstrate tyrosyl hyperfine splitting as marked in (B) by *.

### DOPA is a modification by oxidation of tyrosine residues

**Figure S3.**
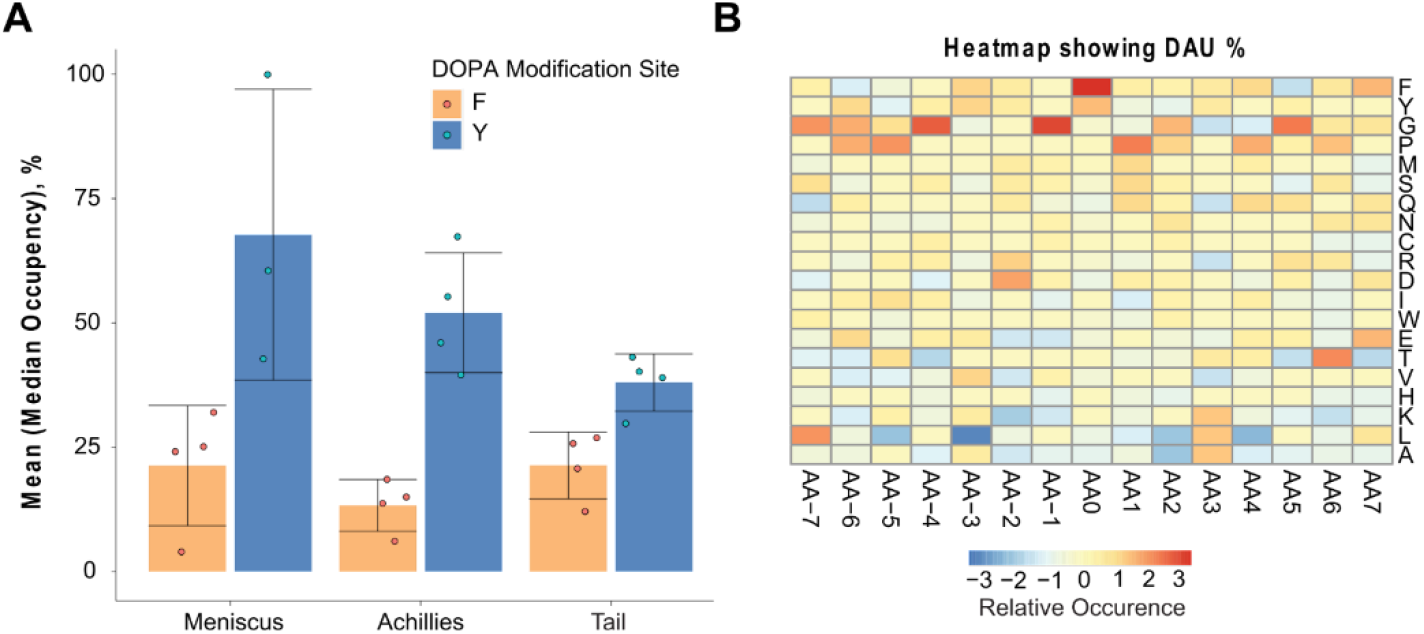
Mass spectrometric analysis of collagen tissues. (**A**) The bar plots (mean ± standard deviation) give the average of median values per modification site (either F or Y) in each sample (for each tissue four biological samples). Occupancy (%) in these peptides was calculated using this formula: 100*Intensity of the Modified Peptide/Sum of intensities of modified and unmodified peptide. (**B**) The motif enrichment was calculated using dagLogo (*29*) using standard settings with only collagenous peptides depicting all amino acid relative occurrence. The central amino acid (AA) position is centered at position 0. The y-axis shows the uni letter code of the AA. The heatmap gives the relative occurrence at any position as differential amino acid usage (DAU, %), which is a relative measurement for the occurrences of a motif (motif enrichment analysis).

### L-DOPA is an efficient radical scavenger

**Figure S4.**
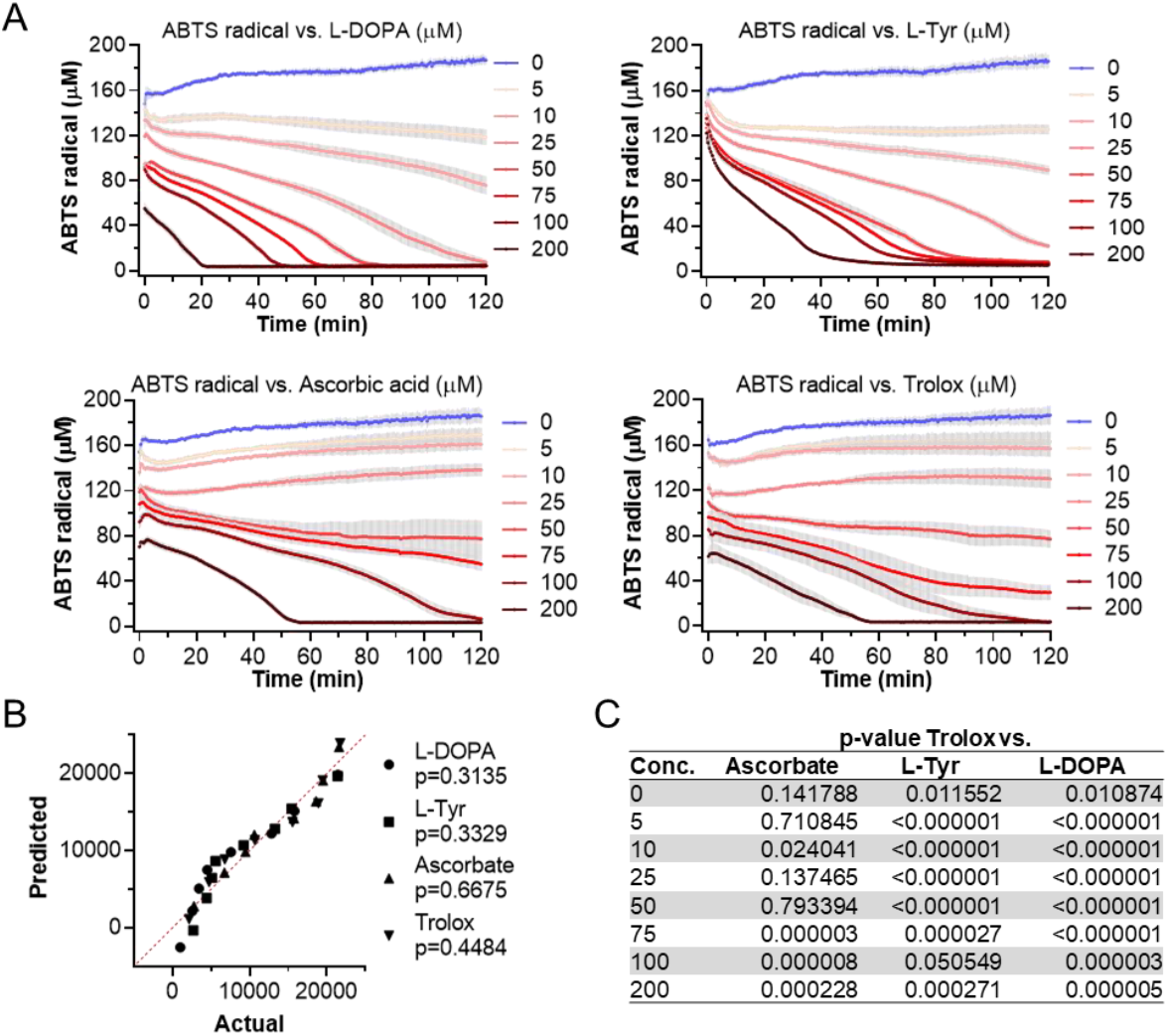
Radical scavenging of amino acids and ascorbic acid in ABTS assay. (**A**) The reaction coordinates of the indicated substances in the ABTS radical scavenging assay were measured as O.D. at λ=734 nm. The O.D. was converted to the ABTS (μM) by the conversion factor y=y*k (k=138.652). Recording in the plate reader started one minute after adding the reactants, so initial scavenging could not be captured. The data represent n=4 independent measurements of each two technical replicates (mean ± SEM). (**B**) Normality of sample population (ABTS area under curve from the mean of four replicates, each measured from two tech. replicates) was tested with the Shapiro-Wilk test (null hypothesis: sample came from a normally distributed population, p≧0.05), here visualized as QQ-plot. p<0.05 indicates significant deviation from normal distribution of the data set. Here, no data set diverged from normal distribution. (**C**) p-values for t-test of Trolox versus the indicated substances. Significance (p<0.05, difference from Trolox) was calculated by multiple t-tests (Holm-Sidak method, no consistent SD, α=0.05).

**Figure S5.**
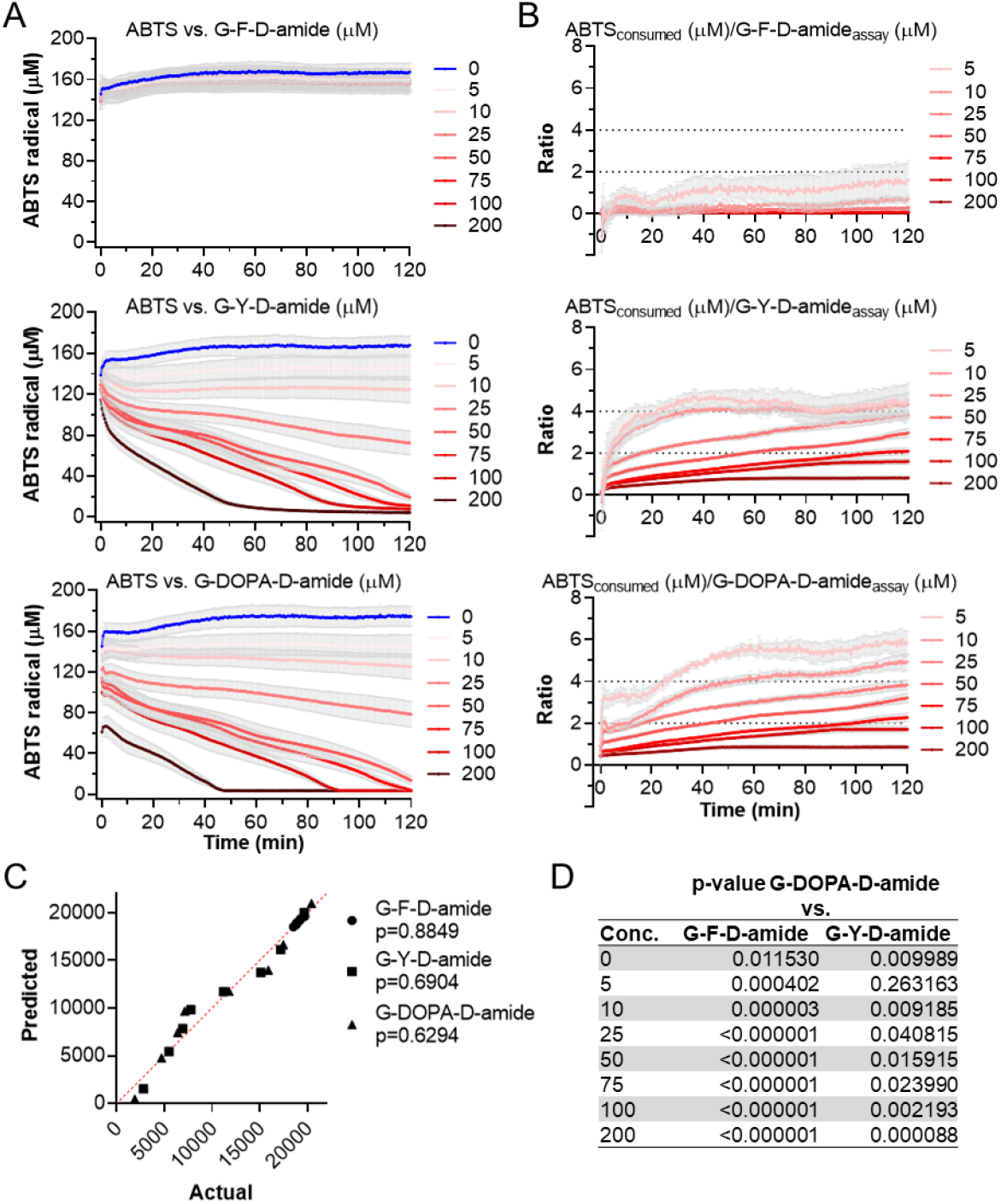
Radical scavenging of tripeptides in ABTS assay. (**A**) ABTS radical concentration over time when scavenged by the indicated tripeptide at different initial concentrations. The O.D. measured at λ=734 nm was converted to the ABTS radical concentration (μM) by the conversion factor y=y*k (k=138.652). The data represent n=4 independent measurements of each two technical replicates (mean ± SEM). (**B**) The corresponding ratios between consumed ABTS (calculated averaging the tech. replicates and subtracting them from the respective c=0 μM) and peptides was calculated by dividing the concentration of consumed radical by the assay concentration of the scavenger. Please note that the apparent radical scavenging of the G-F-D peptide (top) can be attributed to the artifacts when calculating with small values. (**C**) Normality of sample population (ABTS area under curve from the mean of four replicates, each measured from two tech. replicates) was tested with the Shapiro-Wilk test (null hypothesis: sample came from a normally distributed population, p?0.05), here visualized as QQ-plot. p<0.05 indicates significant deviation from normal distribution of the data set. Here, no data set diverged from normal distribution. (**D**) p-values for t-test of G-DOPA-D-amide versus the indicated substances. Significance (p<0.05, difference from G-DOPA-D-amide) was calculated by multiple t-tests (Holm-Sidak method, no consistent SD, α=0.05).

**Figure S6.**
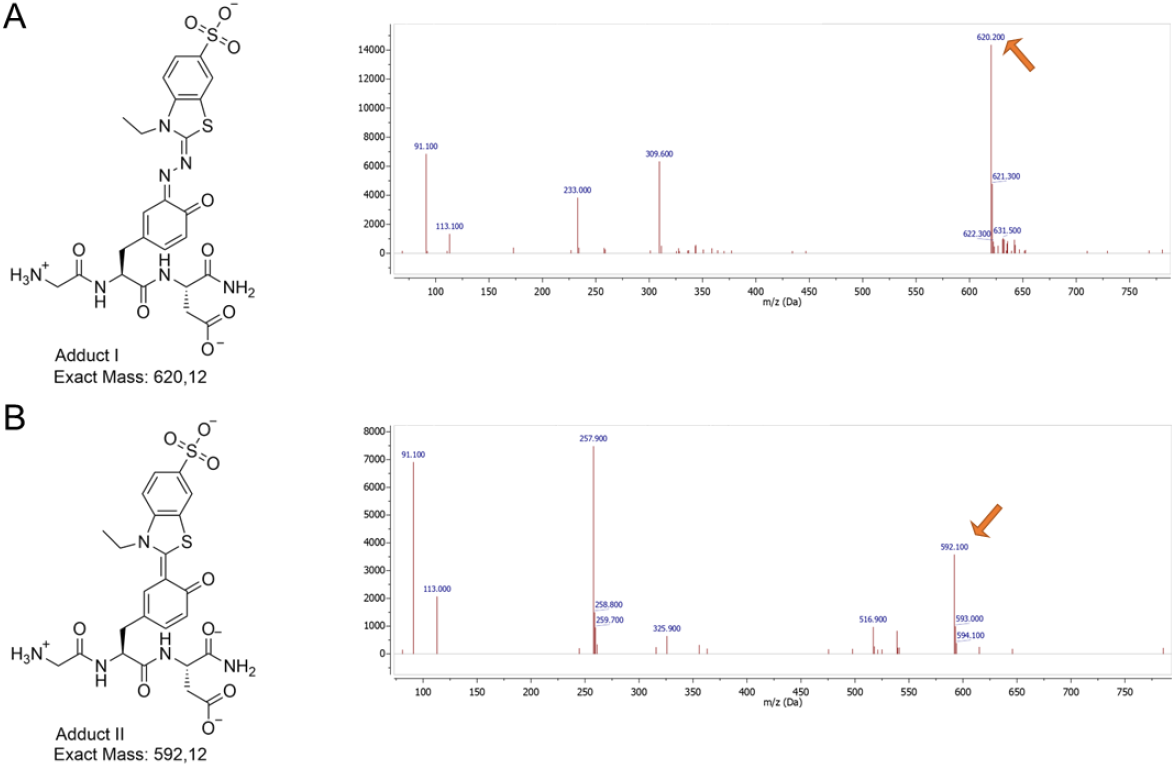
Formation of Tyr-ABTS adduct in ABTS assay. The reaction product was measured by HPLC/MS with ESI. The data was processed with MestReNova (v. 14.2.1) software (see also Experimental section). (**A**) Adduct I between G-Y-D-amide and the ABTS radical fragment “272 m/z” reported earlier (*38, 52*). (**B**) Adduct two between G-Y-D-amide and a possible smaller fragment of 239.98 m/z.

### DOPA radical scavenging produced peroxide

**Figure S7.**
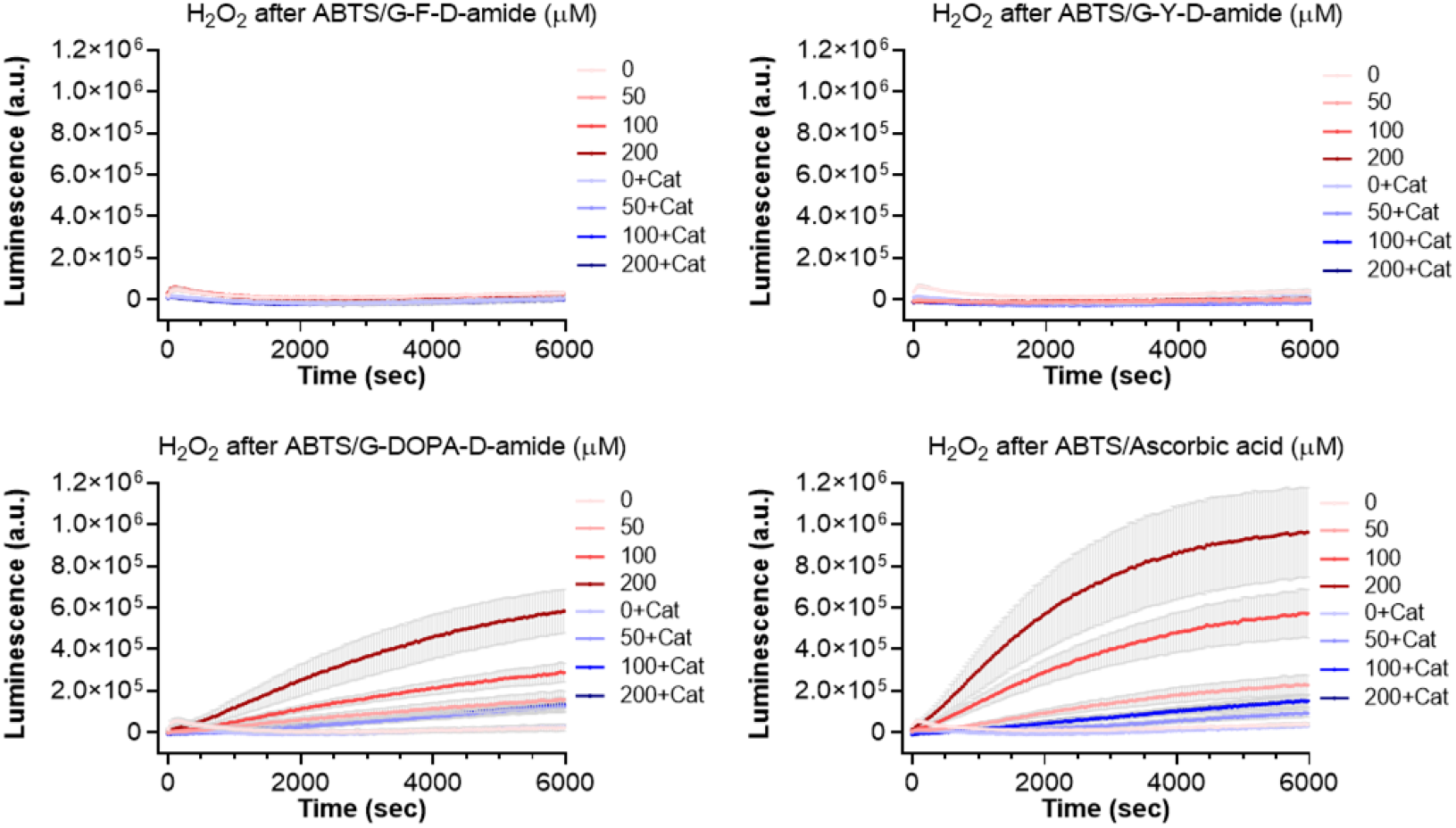
Hydrogen peroxide measurement by luminescence probe. Hydrogen peroxide detection by luminescence of boric acid-based HyPerBlu after ABTS assay. Samples were taken directly from ABTS assay with the named substances in the indicated concentrations and incubated with catalase (Cat) where indicated before detection. Luminescence was recorded for 6000 s at intervals of 42 s (n=4 experiments with each 2 tech. replicates, mean ± SEM). The values were background corrected with the c=0 μM H_2_O_2_ of the corresponding hydrogen peroxide standard.

**Figure S8.**
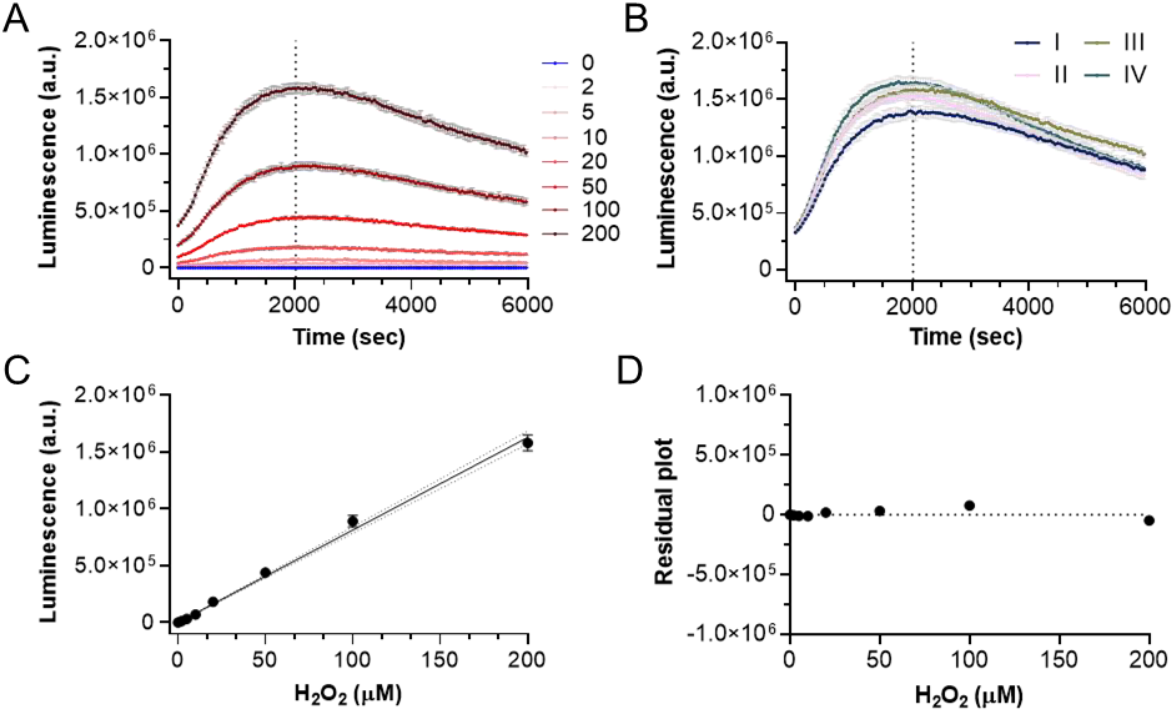
Quantification of hydrogen peroxide measurement by luminescence probe. (**A**) This representative H_2_O_2_ standard (here: repl. III) in the indicated assay concentrations of 0-200 μM was detected by HyPerBlu. The kinetic was recorded for 6000 s at intervals of 42 s (repeated for each of the n=4 replicates with each 2 tech. replicates, mean ± SEM). The control with 0 μM H_2_O_2_ standard served as background control and thus was used for baseline subtraction. For the purpose of orientation, the time of quantification at t=2016 s is indicated as a dashed line. (**B**) Typically, for all four replicates (I-IV) of H_2_O_2_ measurements, the highest concentration of 200 μM peaked at t=~2016 s. For this reason, all four measurements were quantified at this time using the corresponding background corrected values of the luminescence. (**C**) For the interpolation of unknown values at the given time from the H_2_O_2_ standard, the background corrected luminescence values were plotted against the corresponding concentrations (here: repl. III as representative plot). The standard curve (solid line) was fitted as linear regression through y_intercept_=0 (here: y=8143*x) with error given as dashed line representing the 95% confidence interval. (**D**) The residual plot (here: repl. III as representative plot) for the linear regression shown in (C).

**Figure S9.**
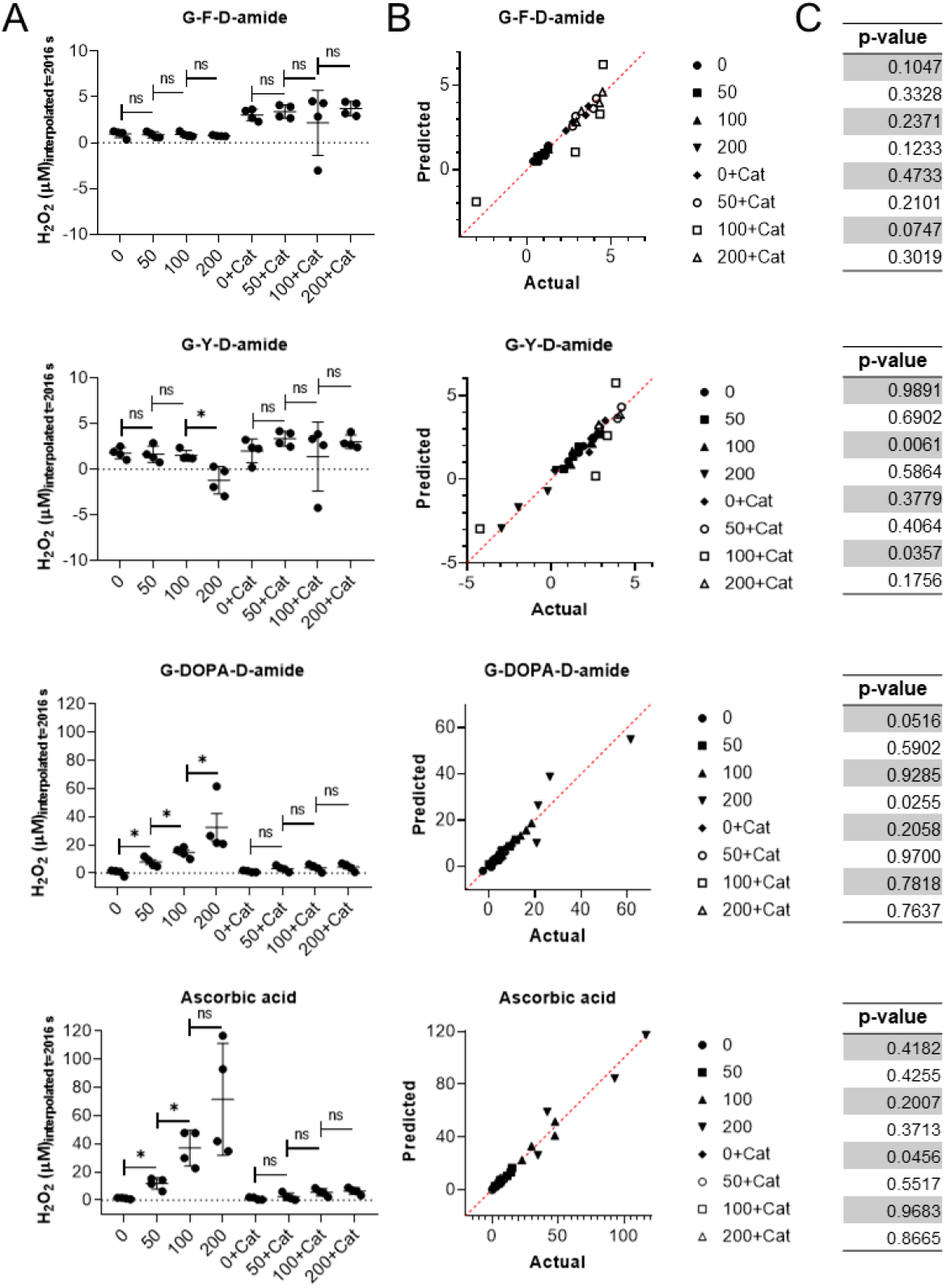
Concentration-dependent hydrogen peroxide formation after ABTS radical quenching by G-DOPA-D-amide and ascorbic acid. The hydrogen peroxide (H_2_O_2_) measurement after ABTS assay (ABTS concentrations indicated on the x-axis) was carried out in the presence and absence of catalase (Cat). (**A**) The significance of the increase in H_2_O_2_ concentration, quantified at t=2016 s (mind the different y-axis scaling for better readability), dependent on the ABTS concentration used was tested by non-parametric Mann-Whitney test (one-sided, confidence level 95%). For G-F-D-amide, we report the p-values with 0.3429, 0.3429, 0.4429, 0.2429, 0.4429, & 0.4429; for G-Y-D-amide with 0.4429, 0.3429, 0.0143, 0.0571,0.2429, & 0.4429; for G-DOPA-D-amide with 0.0143, 0.0286, 0.0143, 0.0571,0.4429, & 0.3429; for ascorbic acid with 0.0143, 0.0143, 0.1714, 0.3429, 0.1000, & 0.2429. Significance of increase is indicated by an asterisk (*, p<0.05) in the graphic, else it is marked as non-significant (ns, p≧0.05). (**B**) Normality of sample population (H_2_O_2_ concentration in the four replicates, each measured from two tech. replicates) was tested with the Shapiro-Wilk test (null hypothesis: sample came from a normally distributed population, p≧0.05), here visualized as QQ-plot. p<0.05 indicates significant deviation from normal distribution of the data set. As single data sets diverged from normal distribution, Mann-Whitney test was chosen for significance test. (**C**) p-values corresponding to the data sets of Shapiro-Wilk normality tests in legend of (B).

**Table S2.** Quantification of hydrogen peroxide in presence of catalase. As control for specificity of the luminescence probe for hydrogen peroxide, catalase was added to the reaction as control. Increase of quantified peroxide concentration in presence and absence of catalase was determined with non-parametric Mann-Whitney test (one-sided, confidence level 95%). This table reports the corresponding p-values. The normality tests for the data sets are reported in Figure S7 with the respective p-values. For visualization, see Figure 5 in the main text).

| Substance | Experiment | p-value |
| --- | --- | --- |
| G-F-D-amide | 0 vs. 0+Cat | 0.0143 |
| G-F-D-amide | 50 vs. 50+Cat | 0.0143 |
| G-F-D-amide | 100 vs. 100+Cat | 0.1714 |
| G-F-D-amide | 200 vs. 200+Cat | 0.0143 |
| G-Y-D-amide | 0 vs. 0+Cat | 0.3429 |
| G-Y-D-amide | 50 vs. 50+Cat | 0.0571 |
| G-Y-D-amide | 100 vs. 100+Cat | 0.1714 |
| G-Y-D-amide | 200 vs. 200+Cat | 0.0143 |
| G-DOPA-D-amide | 0 vs. 0+Cat | 0.4429 |
| G-DOPA-D-amide | 50 vs. 50+Cat | 0.0571 |
| G-DOPA-D-amide | 100 vs. 100+Cat | 0.0143 |
| G-DOPA-D-amide | 200 vs. 200+Cat | 0.0143 |
| Ascorbic acid | 0 vs. 0+Cat | 0.5000 |
| Ascorbic acid | 50 vs. 50+Cat | 0.0143 |
| Ascorbic acid | 100 vs. 100+Cat | 0.0143 |
| Ascorbic acid | 200 vs. 200+Cat | 0.0143 |

**Figure S10.**
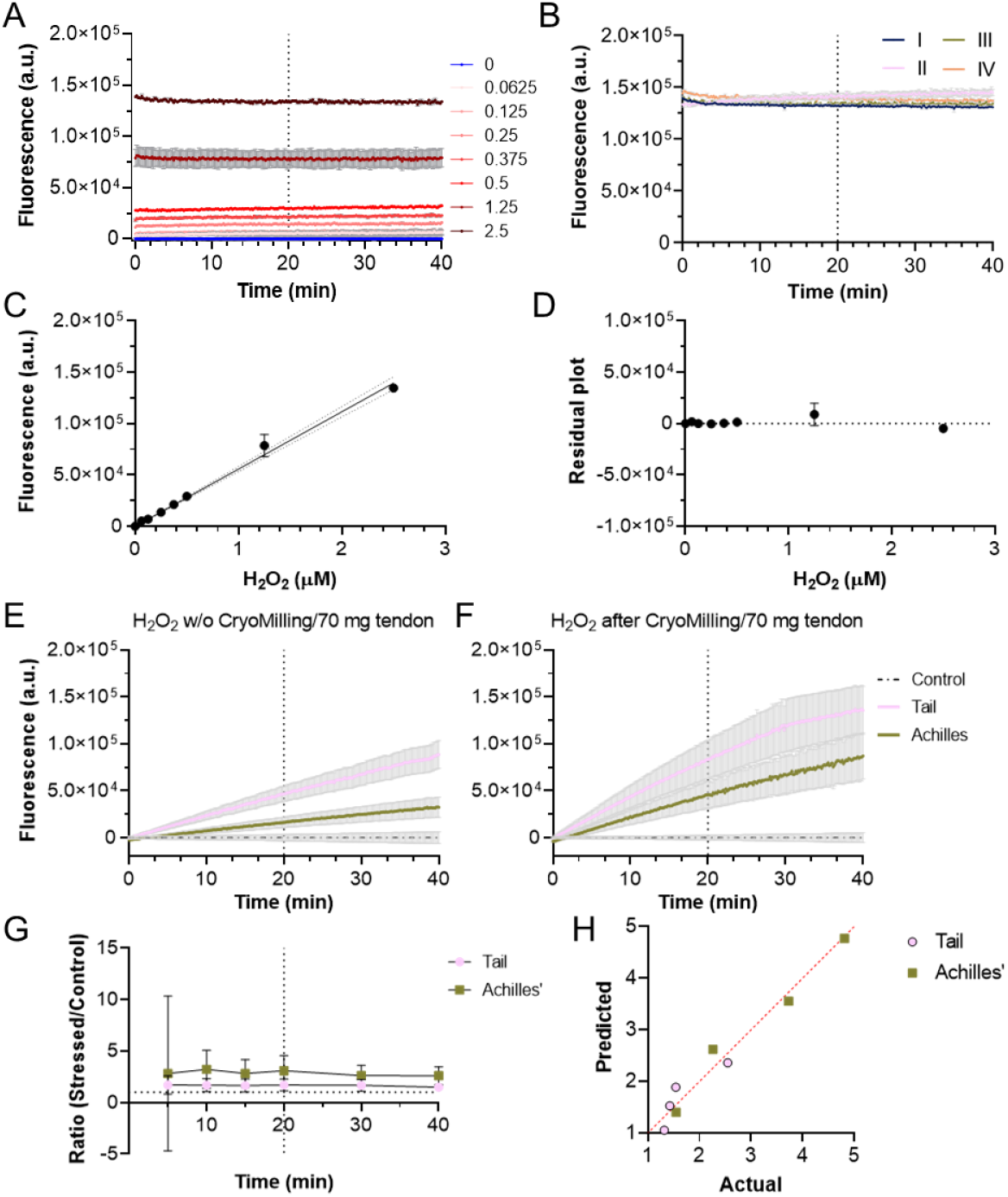
Quantification of hydrogen peroxide measurement by near-infrared horseradish peroxidase assay. (**A**) This representative H_2_O_2_ standard (here: repl. III) in the indicated assay concentrations of 0-2.5 μM was detected by near-infrared horseradish peroxidase kit (Abcam ab138886). The kinetic was recorded for 40 min at intervals of 10 s (repeated for each of the n=4 replicates with each 2 tech. replicates, mean ± SEM). The control with 0 μM H_2_O_2_ standard served as background control and thus was used for baseline subtraction. For orientation, the time of quantification at t=20 min is indicated as a vertical dashed line. (**B**) Typically, for all four replicates (I-IV) of H_2_O_2_ measurements, the fluorescence signal (here at highest concentration of 2.5 μM) remained relatively stable during the whole measurement time of 40 min. For this reason, all three measurements were quantified at half-time. (**C**) For the interpolation of unknown values at the given time from the H_2_O_2_ standard, the background corrected fluorescence values were plotted against the corresponding concentrations (here: repl. III as representative plot). The standard curve (solid line) was fitted as linear regression through y_intercept_=0 (here: y=55757*x) with error given as dashed line representing the 95% confidence interval. (D) The residual plot (here: repl. III as representative plot) for the linear regression shown in (C). (**E**) The graphic shows the kinetics of untreated tissue samples (~70 mg tissue/2 zirconium oxide marbles) and the background control (dashed line). As background control, an empty reaction tube with two zirconium oxide marbles was used to carry out the assay reaction in the same way as for the tissue samples; the fluorescence values were used for baseline subtraction (repeated for each of the n=4 replicates with each 2 tech. replicates, mean ± SEM). (**F**) The graphic shows the kinetics of cryomilled tissue samples (~70 mg tissue/2 zirconium oxide marbles) and the background control (dashed line). As background control, an empty reaction tube with two zirconium oxide marbles was used to carry out the assay reaction in the same way as for the tissue samples; the fluorescence values were used for baseline subtraction (repeated for each of the n=4 replicates with each 2 tech. replicates, mean ± SEM). (**G**) While the absolute values for the fluorescence values increased over time and thus also the corresponding quantification for c(H_2_O_2_), we decided to use the ratio cyromilled (=stressed) to untreated (=control) fluorescence values instead. As indicated, the background corrected values retrieved at t=5 min, 10 min, 15 min, 20 min, 30 min, and 40 min remained stable after ~15 min measurement time and stayed constant until the end of the assay time. (**H**) Shapiro-Wilk test for normality (null hypothesis: sample came from a normally distributed population, p≧0.05) for tail and Achilles’ tendon ratios (stressed/control) at t=20 min (QQ plot). For tail samples, p-value was reported with 0.0613, for Achilles with 0.7867.

